# Allosteric disulfide control of ligand binding and endocytosis of KIR2DL4, the natural killer cell receptor for HLA-G

**DOI:** 10.64898/2026.02.18.706601

**Authors:** Sumati Rajagopalan, Joyce Chiu, Priyanka Chaurasia, Jinghua Lu, George M. Mastorakos, Saurav Majumder, Kristof Nolan, Jan Petersen, Erin J. Adams, Peter D. Sun, Jamie Rossjohn, Philip J. Hogg, Eric O. Long

**Affiliations:** Molecular and Cellular Immunology Section, Laboratory of Immunogenetics, National Institute of Allergy and Infectious Diseases, National Institutes of Health, Rockville, MD 20852, USA; Haematology Research Unit, School of Clinical Medicine, University of New South Wales, Sydney, NSW, Australia; Infection and Immunity Program and Department of Biochemistry and Molecular Biology, Biomedicine Discovery Institute, Monash University, Clayton, Victoria 3800, Australia; Structural Immunology Section, Laboratory of Immunogenetics, National Institute of Allergy and Infectious Diseases, National Institutes of Health, Rockville, MD 20852, USA; Department of Biochemistry and Molecular Biology, University of Chicago, Chicago, IL 60637; Centenary Institute, University of Sydney, Camperdown, NSW, Australia; School of Life Sciences, University of Technology Sydney, Sydney NSW 2007, Australia; Scienza Health, Inc; Center for Blood Oxygen Transport and Hemostasis, University of Maryland School of Medicine, Baltimore, MD 21201, USA; AstraZeneca, Inc

**Keywords:** reproduction, fetal development, decidua, trophoblast, HLA-G, natural killer cell, allosteric disulfide, KIR2DL4, mutagenesis

## Abstract

Human Leukocyte Antigen (HLA)-G is selectively expressed by fetal trophoblast cells that invade maternal tissue and encounter maternal natural killer (NK) cells early in pregnancy. In NK cells, the endosomal receptor KIR2DL4 responds to soluble HLA-G by inducing a broad transcriptional program to support placental development. Structural features of KIR2DL4 that control ligand binding and endocytosis are unclear. Random mutagenesis revealed that three cysteines in the first immunoglobulin domain of KIR2DL4 regulate endocytosis and uptake of HLA-G. We found that the atypical Cys10-Cys28 disulfide bond observed in the KIR2DL4 crystal structure is an allosteric disulfide with potential to switch to a conventional Cys28-Cys74 bond. KIR2DL4 in human cells exists in both disulfide-bonded states, as shown by mass spectrometry analysis. The Cys10-Cys28 bond in a Cys74Ser KIR2DL4 mutant was reduced by protein disulfide isomerase (PDI) in vitro. Inhibition of PDI prevented HLA-G uptake by KIR2DL4 in both transfected 293T cells and primary NK cells. Mutants in the Cys10-Cys28 configuration endocytosed spontaneously but did not bind HLA-G. Conversely, KIR2DL4 with a Cys28-Cys74 bond was at the plasma membrane and responded to soluble HLA-G by endocytosis and transcription of the interferon stimulated gene *IFI44L*, like wild-type cells. A purified Cys10Leu KIR2DL4 mutant, which exhibited a reduced tendency to oligomerize, bound HLA-G in a peptide dependent manner with an affinity in the low micromolar range. Disulfide switching from the Cys10-Cys28 to the Cys28-Cys74 form correlated with a distant allosteric change, as predicted by AlphaFold, that could reorient a D0 domain loop to facilitate HLA-G binding. Thus, conversion from an inactive state to an HLA-G binding form regulates KIR2DL4 cellular localization and function to promote fetal development.

## Introduction

Human natural killer (NK) cells play an important role in immune defense and reproduction (*1–3*). They respond to alterations in healthy cells caused by cellular stress, infection, and malignancy by killing these aberrant cells and producing cytokines and chemokines (*2*). In the context of pregnancy, NK cells, the most abundant lymphocytes at the maternal-fetal interface, are not killers. Instead, they play a constructive role by interacting with and responding to fetal trophoblast cells. Data suggest that a secretory response by NK cells promotes remodeling of the maternal vasculature to support a successful pregnancy (*4–6*).

NK cells express an array of receptors to sense their environment. Integration of signals from both activatory and inhibitory receptors that interact with ligands on target cells determines NK cell responses (*7*). The killer-cell Ig-like receptors (KIR), most of which recognize HLA class I molecules, regulate NK cell activation through expression of both inhibitory and activatory KIR family members. KIR2DL4 is a unique member of the KIR family whose expression, unlike other KIR, occurs in most NK cells and is higher in decidual NK cells in early pregnancy (*5, 8*). KIR2DL4 has two Ig-like domains (termed D0 and D2) and activates a secretory response after engagement by a soluble agonist antibody or by soluble HLA-G (*9*). KIR2DL4 resides mainly in early endosomes and binds soluble HLA-G to stimulate a broad transcriptional response, including an interferon stimulated gene (ISG) response in primary NK cells that may protect NK cells from pathogens (*10*). These responses are relevant in the context of early pregnancy where HLA-G expression by fetal trophoblast cells is controlled spatially and temporally as they invade the maternal decidua following implantation of the embryo (*5*).

Despite evidence of an endosomal signaling pathway initiated by a KIR2DL4–HLA-G interaction (*11*), direct binding of HLA-G to KIR2DL4 protein has not been reported, and previous findings showed no biomolecular interaction (*12*). Even though KIR2DL4 is required for responses to HLA-G and an agonist antibody to KIR2DL4 is sufficient for activation (*13, 14*), the requirement for a coreceptor for HLA-G mediated activation of NK cells was not excluded. We recently reported a strong correlation in the transcriptional response of primary NK cells stimulated with soluble HLA-G with their response to the agonist monoclonal antibody #33 (mAb #33), consistent with direct engagement of KIR2DL4 with HLA-G (*10*). To examine how KIR2DL4 interacts with HLA-G, we chose a random mutagenesis approach and mapped residues in the extracellular domain that are required for a physical and functional interaction with HLA-G. Several independent mutations implicated three cysteine residues in the D0 domain that controlled binding of HLA-G and endocytosis of KIR2DL4. We provide evidence for an allosteric disulfide switch between these three cysteines that regulates both endocytosis and ligand binding. Consistent with this, we show that a purified cysteine mutant of KIR2DL4 in the functional Cys28-Cys74 configuration does in fact bind HLA-G in vitro in a peptide-dependent manner.

## Results

### Random mutagenesis of KIR2DL4 to map its interaction with HLA-G

Random mutagenesis of a cDNA fragment that encodes the KIR2DL4 extracellular domain was performed using a low-fidelity PCR reaction calibrated to generate approximately 1 to 2 base change per 1000 bp (Fig. S1A). HA-tagged KIR2DL4 with single or double amino acid changes (n=293) were transfected individually in 293T cells and examined for their ability to bind HLA-G (Fig. 1A and S1B). Transfected cells were incubated for 2 hours at 37°C with Alexa-594 coupled to an anti-HA tag antibody and with soluble HLA-G coupled to Alexa-647 (Fig. S1B). HLA-G had been refolded with the single peptide KGPPAALTL, which was the most abundant peptide eluted from HLA-G obtained from placenta (*15, 16*). The distribution of KIR2DL4 and of HLA-G was determined by confocal microscopy (Fig. 1B). As shown previously (*14*), soluble HLA-G was endocytosed along with KIR2DL4 into endosomes. As controls, soluble HLA-E coupled to Alexa-647 was not bound nor internalized into KIR2DL4**^+^** endosomes, and HLA-G-647 was not internalized in 293T cells expressing an HA-tagged 2B4 (*CD244*) receptor (Fig. 1B). Out of 216 clones with interpretable images of distribution at the plasma membrane or in intracellular vesicular compartments (vesicles), 153 (71%) had a phenotype that resembled wild-type KIR2DL4. The remaining 63 mutants had distinct non-wild-type phenotypes (Fig. 1C).

**Figure 1:**
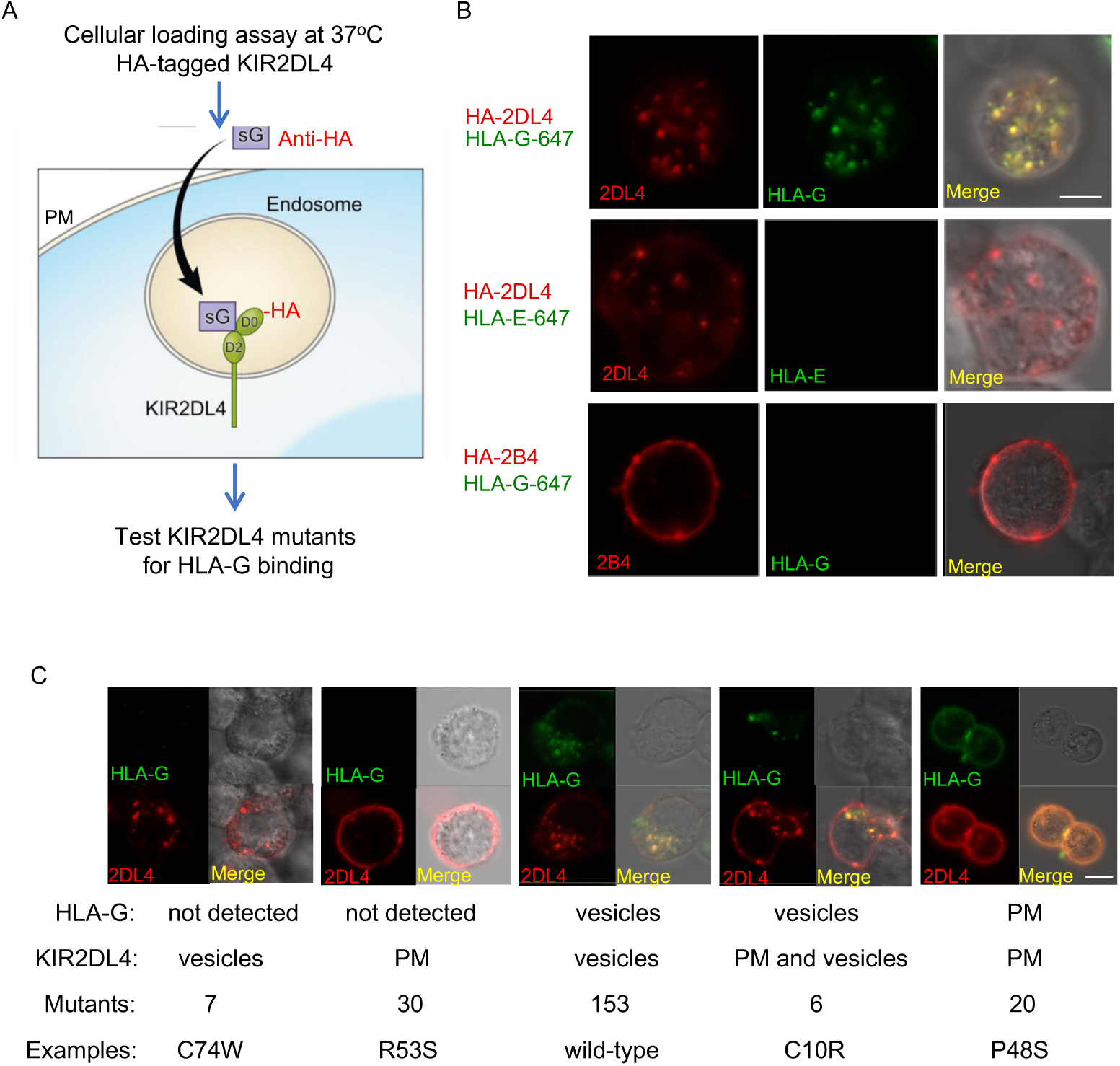
Random mutagenesis screen. (**A**) Soluble HLA-G (sG) is internalized with HA-tagged KIR2DL4 into endosomes, as shown by incubation for 2 h at 37°C with anti-HA coupled to Alexa-594 and soluble HLA-G coupled to Alexa-647. The loading assay was performed with 293T cells transfected with HA-tagged KIR2DL4. (**B**) Confocal microscopy showing the location of internalized KIR2DL4 (*red*) and soluble HLA-G (*green*). An overlay is shown in the right panel (*Merge*). As control, HLA-G was replaced by HLA-E, also coupled to Alexa-647 (*middle panels*). 293T cells transfected with HA-tagged receptor 2B4 were also incubated with HLA-G coupled to Alexa-647 (*bottom panels*). 2B4 (*CD244*) resides at the plasma membrane. (**C**) The loading assay was used to screen KIR2DL4 mutants and test for HLA-G binding. The main parameter measured was presence or absence of HLA-G and location of detectable HLA-G. Confocal images representative of the five categories of mutants are shown. A DIC image is shown in the upper right panel. The parameters indicated under the images include HLA-G presence and location, location of KIR2DL4, the number of mutants in each of the five categories, and an example of a mutation for each category.

The main objective was to identify KIR2DL4 mutants that did not bind HLA-G and were otherwise in vesicles, similar to wild-type KIR2DL4 in the absence of HLA-G (*14*). Only 7 mutants (out of 63) had that phenotype (Fig. 1C, *first panel*, Dataset S1). The more common phenotype of KIR2DL4 mutants that did not bind HLA-G (30 out of 37) was an even distribution at the plasma membrane (Fig. 1C, *second panel*). Mutations with that phenotype were distributed throughout the D0 and D2 Ig domains (Dataset S1), consistent with internalization that is controlled by the KIR2DL4 extracellular domain (*17*).

Two distribution patterns were observed among the 26 mutants that bound HLA-G. Six mutants displayed a hybrid phenotype: colocalization of HLA-G with KIR2DL4 in vesicles, like wild-type, but with some KIR2DL4 remaining the plasma membrane (Fig. 1C, *fourth panel*), suggesting that internalization of these 6 mutants was facilitated by HLA-G. To test this, the same mutants were examined in the absence of HLA-G. Results showed that, indeed, they remained at the plasma membrane (Fig. S1C). The other 20 mutants were detected at the plasma membrane along with HLA-G (Fig. 1C, *right most panel*). Overall, these results showed that 50 out of the 63 mutants that had a non-wild-type phenotype (Dataset S1) failed to internalize, reinforcing the evidence that KIR2DL4 internalization is controlled by the extracellular domain (*17*).

The KIR2DL4 D0 domain includes 3 cysteines at positions 10, 28, and 74. Among the 7 mutants that did not bind HLA-G but resided in vesicles, two included a mutation at Cys74, such as C74W (Fig. 1C). This suggested that KIR2DL4 with a Cys10-Cys28 disulfide bond, as seen in the KIR2DL4 crystal structure (*12*), does not bind HLA-G but is otherwise distributed in vesicles like wild-type KIR2DL4 (*14*). Conversely, out of the 6 mutants with a hybrid phenotype, namely distribution at the plasma membrane and in vesicles but colocalization with HLA-G in vesicles only, one had a mutation at Cys10 (C10R). This observation showed that KIR2DL4 with the canonical Ig domain disulfide bond Cys28-Cys74 is detectable at the plasma membrane in the absence of HLA-G (Fig. S1C, left). This C10R mutant was at the plasma membrane only, demonstrating that its internalization was facilitated by soluble HLA-G. Conversely, we tested if KIR2DL4 mutants retained at the plasma membrane with bound HLA-G, such as P48S (Fig. 1C), could internalize on their own. In the absence of HLA-G, mutant P48S was in vesicles (Fig. S1C, *right*), showing that, in this case, HLA-G binding caused retention at the plasma membrane.

Several targeted mutations were generated to confirm these results. The phenotypes of mutants C74W and C10R (Fig. 1C) were reproduced with targeted mutations C74A and C10S, respectively (Fig. S1D, and Table 1). These mutations also supported the conclusion that KIR2DL4 with a Cys28-Cys74 bond is detectable at the plasma membrane in the absence of HLA-G. Conversely, HLA-G prevented the internalization of some KIR2DL4 mutants, such as P48S, which otherwise endocytosed spontaneously (Fig. S1C).

**Table 1:** Cysteine mutants in the D0 domain of KIR2DL4 and their impact on the distribution of KIR2DL4 and its interaction with HLA-G. V = vesicular; PM = Plasma membrane.

| KIR2DL4<br>Cys<br>mutants | KIR2DL4<br>location | KIR2DL4<br>location upon<br>sHLA-G load | sHLA-G<br>location | Disulfide Bonds<br>in D0 domain |
| --- | --- | --- | --- | --- |
| WT | V | V | V | 3 possible |
| C10S | PM | PM and V | V | C28-C74 |
| C10A | PM | PM and V | V |  |
| C10R | PM | PM and V | V |  |
| C28S | PM | PM | None | C10-C74 |
| C28A | PM | PM | None |  |
| C28R | PM | PM | None |  |
| C74S | V | V | Few V | C10-C28 |
| C74A | V | V | None |  |
| C74R | V | V | Few V |  |
| C74W | V | V | None |  |

### An allosteric disulfide bond in KIR2DL4

The impact of cysteine mutations on the localization of KIR2DL4 led us to compare its sequence with that of other D0 domain found in KIR2DL5 and the KIR3DL family (Fig. 2A). The cysteine at position 10 of KIR2DL4 is absent in these other KIRs, which have a conserved leucine at that position. The single disulfide bond in those D0 domains is a canonical Ig domain *trans*-beta sheet bond (*18*), comparable to the Cys28-Cys79 bond in the D1 Ig domain of KIR2DL1 (Fig. 2A, B). Of the three potential disulfide bonds (Cys10-Cys28, Cys28-Cys74, and Cys10-Cys74), the one visible in the KIR2DL4 crystal structure (*12*) is Cys10-Cys28, leaving a free Cys at position 74 (Fig. 2C).

**Figure 2:**
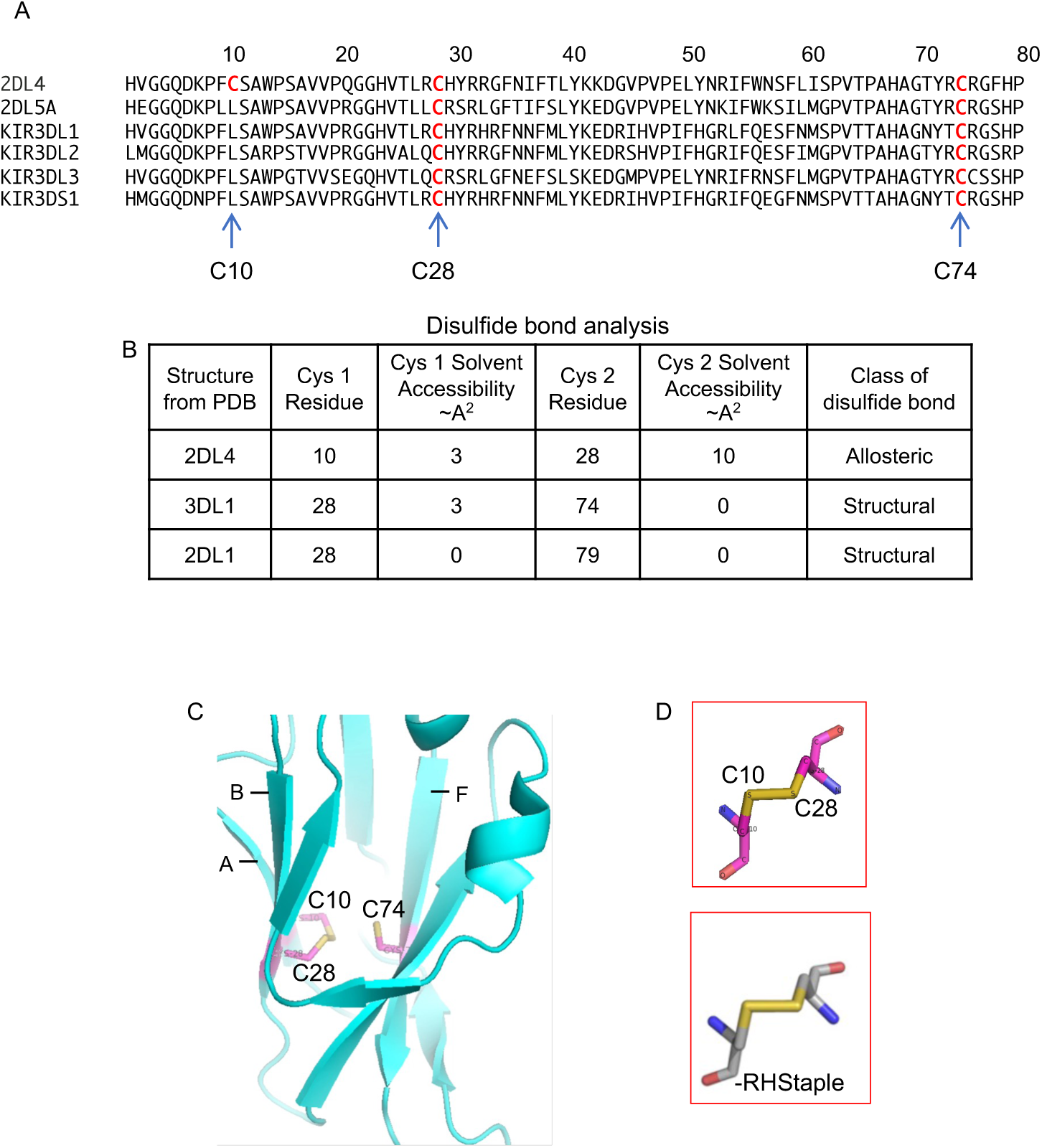
A stressed disulfide bond typical of an allosteric bond in the crystal structure of KIR2DL4. (**A**) The KIR2DL4 D0 domain has a third Cys residue not found in other D0 domains. Sequence alignment of D0 domains with Cys residues indicated by an arrow and shown in red (**B**) Solvent accessibility of disulfide bonds visible in crystal structures of the first Ig domain of KIRs was determined by DSSP (define secondary structure of proteins) algorithm. The first Ig domain of KIR2DL4 and KIR3DL1 is a D0 domain, whereas the first Ig domain of KIR2DL4 is a D1 domain. (**C**) Position of the three Cys residues in the crystal structure of KIR2DL4 (3WYR). Cys10, Cys28 and Cys74 are on the A strand, B strand and F strands, respectively, are as indicated. (**D**) The disulfide bond configuration in the KIR2DL4 D0 domain is an –RHstaple.

Disulfide bonds that contribute to protein folding and stabilization tend to be inert and are referred to as structural bonds (*18*). Labile disulfide bonds that can switch to a reduced form and back to the oxidized bond regulate protein function or localization and are referred to as allosteric bonds (*18*). Such bonds may be stressed and prone to cleavage (*19*). Formation of disulfide bonds may be reversed in reducing conditions or by the enzymatic activity of protein disulfide isomerases (PDI). Examination of the Cys10-Cys28 bond in the D0 domain of KIR2DL4, visible in the crystal structure (*12*), revealed that the second cysteine, Cys28, was partially exposed to solvent (∼10A^2^) (Fig. 2B), compatible with a propensity for thiol disulfide exchange mediated by a protein disulfide isomerase. By comparison, the disulfide bond in the KIR3DL1 D0 domain (Fig. 2B) has low solvent accessibility (∼3A^2^), according to the crystal structure (*20*). Given that KIR2DL4 may be regulated by an allosteric disulfide bond, we examined the configuration of the Cys10-Cys28 disulfide (Fig. 2C). Among the 20 possible disulfide bond conformations, as determined by the geometry of the five dihedral angles that describe the cysteine residue, three of them are characteristic of allosteric bonds (*18, 21*). The Cys10-Cys28 bond in the KIR2DL4 structure has an archetypal –RHstaple allosteric conformation (Fig. 2D), which is characterized by high tensile pre-stress due to stretching of the S-S bond and neighboring α angles (*22*).

### Analysis of Disulfide bonds in KIR2DL4

As cysteine mutations in the D0 domain of KIR2DL4 had an impact on internalization and cellular distribution, we determined what disulfide bond out of the three possible (Fig. 3A) existed in recombinant KIR2DL4 produced in insect cells. Cysteine pairings for disulfide bonds in recombinant KIR2DL4 were mapped using disulfide-linked peptides and mass spectrometry. The conserved Cys123-Cys172 bond in domain D2 (Fig. 3A) served as a reference. In wild-type KIR2DL4, peptides linked by disulfides Cys10-Cys28, Cys10-Cys74, Cys28-Cys74 from D0 and Cys123-Cys172 from D2 were identified and quantified (Fig. 3 B; Fig. S2). Cys10-Cys28 was the more abundant bond in D0 and, as expected, was abundant in the C74S mutant and absent in the C10S mutant (Fig. 3B). The recovery of the reference Cys123-Cys172 bond in the D2 domain was lower (approximately one third of those in the D0 domain) but, as expected, its abundance was virtually identical in the 3 isoforms of KIR2DL4 (wild-type, C10S, and C74S). The Cys28-Cys74 bond in D0 was detected at a lower abundance than Cys10-Cys28, and as expected, was more abundant in the C10S mutant (Fig. 3B). The Cys10-Cys74 bond was detected at a low abundance and was completely absent in the C10S and the C74S mutants (Fig. 3B). In conclusion, wild-type purified KIR2DL4 existed in both configurations, Cys10-Cys28 and Cys28-Cys74, representing independent isoforms distinguished by their disulfide bond in the D0 domain Fig. 3B).

**Figure 3:**
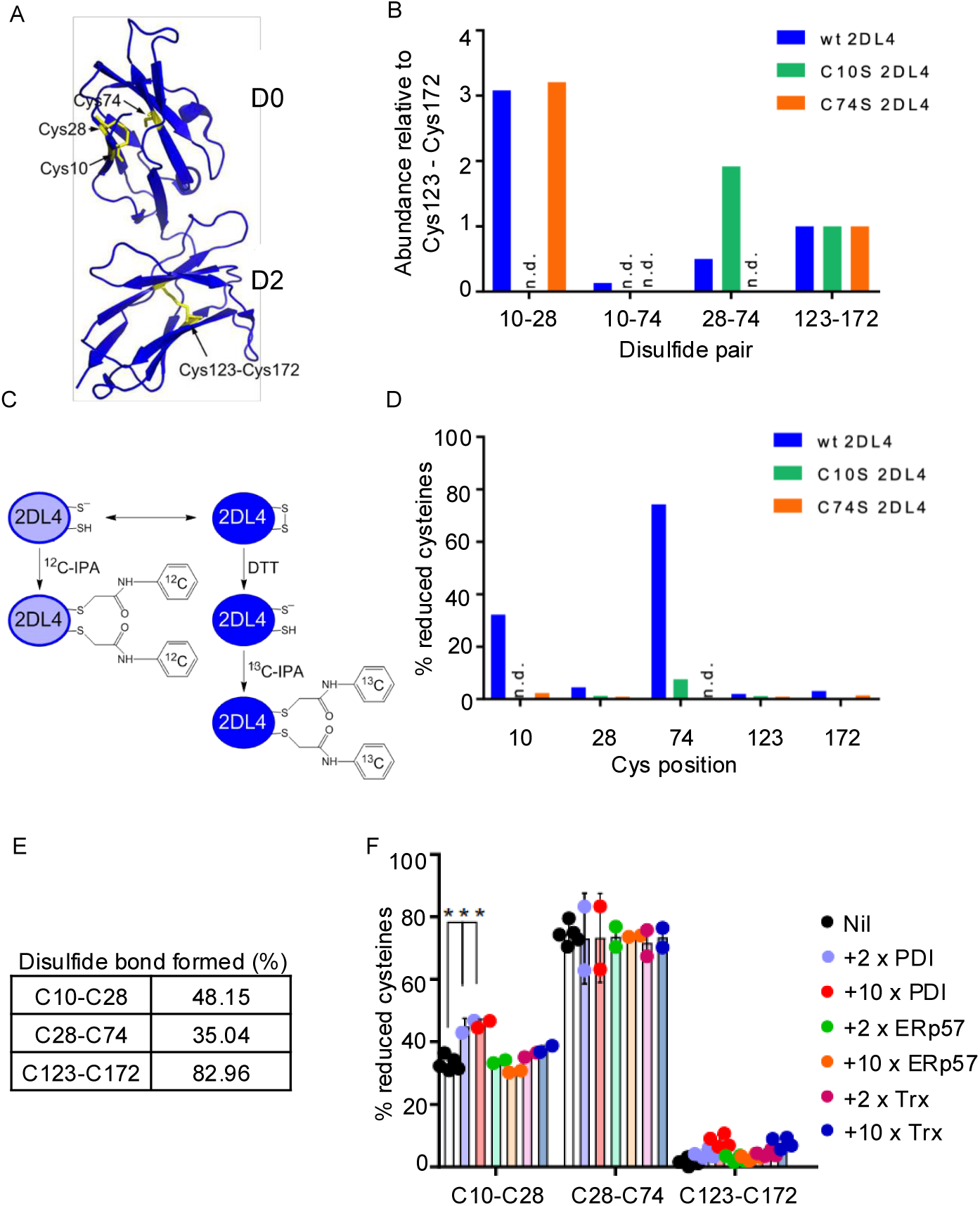
Detection and quantification of disulfide bonds in KIR2DL4 isolated from cells. (**A**) Disulfide bonds in KIR2DL4. The Cys10-Cys28 disulfide bond and the unpaired Cys74 in the D0 domain and the stable Cys123-Cys172 disulfide bond in the D2 domain are shown in yellow in the ribbon structure of KIR2DL4. PDB identifier is 3WYR. (**B**) Quantification of disulfide-linked peptides in recombinant KIR2DL4 by mass spectrometry. Peptide abundance was expressed relative to the peptide linked by Cys123 and Cys172 in the D2 domain. (**C**) Protocol to determine the redox state of KIR2DL4 disulfide bonds, as measured by differential cysteine alkylation and mass spectrometry. Reduced disulfide bond cysteines in KIR2DL4 were alkylated with ^12^C-IPA and the oxidized cysteine thiols with ^13^C-IPA following reduction with DTT. The ratio of ^12^C-IPA to ^13^C-IPA alkylation represents the proportion of the disulfide bonds in the population that are in the reduced state. (**D**) Quantification of cysteine redox states in recombinant KIR2DL4 by differential cysteine alkylation and mass spectrometry. **(E)** Human cell-derived KIR2DL4 was found in both Cys10-Cys28 and Cys28-Cys74 redox states. The redox state of disulfide Cys123-Cys172 is shown for comparison. (**F**) Incubation of purified KIR2DL4 with 3 different protein disulfide isomerases. Reduction of the Cys10-Cys28 bond occurred with PDI at a PDI:KIR2DL4 molar ratio of 2 and 10. The redox state of cysteines were quantified by differential cysteine labelling and mass spectrometry. The error bars (SD) were derived from measurements of 2 to 4 peptides. ***<0.005 as assessed by unpaired, two-tailed Students t-test.

The redox state of cysteines in KIR2DL4 purified from insect cells was then determined by differential cysteine alkylation and mass spectrometry (Fig. 3C; Table S1). Unpaired or reduced cysteines were first labelled with the cysteine alkylator ^12^C-IPA. Disulfide bonds were then reduced by dithiothreitol followed by cysteine alkylation with ^13^C-IPA. The fraction of reduced disulfide bonds was measured from the relative ion abundance of peptides containing ^12^C-IPA or ^13^C-IPA (Fig. 3C). By this approach, the relative abundance of reduced cysteines in wild-type KIR2DL4 was 32.3%, 4.5%, and 74.3% for Cys10, Cys28, and Cy74 respectively (Fig. 3D). Based on these measurements, the calculated abundance of Cys10-Cys28 and Cys28-Cys74 was 67.7% and 25.7%, respectively. This data demonstrated that these two disulfide forms exist in the D0 domain.

Next, we tested if the Cys10-Cys28 and Cys28-Cys74 disulfide pairs existed in KIR2DL4 obtained from human cells, using the same protocol as in Fig. 3C and 3D with the following modifications. Unpaired cysteines on intact 293T-2DL4-gfp cells were first labelled with ^12^C-IPA, followed by cell lysis, pull-down of KIR2DL4 via the GFP tag, and a second labelling with ^12^C-IPA prior to elution. Eluted proteins were separated by gel electrophoresis and those with an approximate mass of 75 kD were isolated. Disulfide bonds were then reduced and the new unpaired cysteines alkylated, this time with ^13^C-IPA (Fig. 3C). The redox states of cysteines were quantified by mass spectrometry analysis. The fraction of reduced disulfide bonds was calculated from the relative ion abundance of peptides containing ^12^C-IPA and ^13^C-IPA (Fig. 3C). The Cys10-Cys28 and the Cys28-Cys74 forms of KIR2DL4 were detected at a 48.15% and 35.04% relative abundance, respectively (Fig. 3E). A KIR2DL4 in the Cys10-Cys74 configuration was not detected, consistent with the very low abundance of reduced Cys28 in recombinant KIR2DL4 (Fig. 3D). The structural Cys123-Cys172 bond in the D2 Ig domain, was present at an 82.96% relative bonded state, equivalent to the total bonded state of the D0 domain (48% + 35%). We concluded that wild-type KIR2DL4 expressed in human cells exists in both disulfide forms (Cys10-Cys28 and Cys28-Cys74), supporting the possibility of an allosteric transition between the two.

### Protein disulfide isomerase controls endocytosis of KIR2DL4

Given that the disulfide-bonded Cys28 in the KIR2DL4 crystal structure is partially exposed to solvent (Fig. 2B), we tested if it was sensitive to cleavage by thiol isomerases. Purified recombinant KIR2DL4 isolated from insect cells was incubated with the three reduced thiol isomerases PDIA1 (*P4HB*), Erp57 (*PDIA3*), and thioredoxin (*TXN*). Major reduction of the Cys10-Cys28 bond but not the Cys28-Cys74 bond was observed in the presence of PDI, but not the other two isomerases (Fig. 3F). Therefore, reduction of Cys10-Cys28 by PDI revealed a selectivity at two levels. One is that the Cys28-Cys74 was resistant, suggesting that the Cys10-Cys28 bond is more accessible to PDI. Second, the lack of reduction by the other two isomerases, Erp57 and Thioredoxin, suggested that accessibility of the Cys10-Cys28 bond to these isomerases is limited.

So far, our results, including data obtained with an unbiased mutagenesis screen, pointed to a major role of disulfide bonds in the D0 domain of KIR2DL4 for trafficking and to their regulation by the activity of protein disulfide isomerase. Several PDI inhibitors were used to examine the cellular distribution of HA-tagged KIR2DL4 in 293T cells incubated with soluble HLA-G coupled to a fluorophore. Two types of inhibitors were used. The first consisted of two thiol blockers that interact irreversibly with free and exposed thiol groups (DTNB and pCMPS). In untreated cells, KIR2DL4 was found mostly in endocytic vesicles together with soluble HLA-G (Fig. 1B and Fig. 4 A-D). In transfected 293T cells that express a high level of KIR2DL4, some KIR2DL4 was detected at the plasma membrane. With increasing concentrations of DTNB (Fig. 4A) or pCMPS (Fig. 4B), the proportion of cells exhibiting KIR2DL4 at the plasma membrane increased. Uptake of HLA-G was sensitive to DTNB and, to a lesser extent, with pCMPS. Incubation with Rutin (quercetin-3-rutinoside), a selective inhibitor of PDI at a concentration that does not inhibit other extracellular thiol isomerases present in the vasculature (*23*) had a similar effect as thiol blockade (Fig. 4C). Blocking of PDI with a specific antibody inhibited internalization of KIR2DL4 but not as efficiently as thiol blockers (Fig. 4D). Thus, blocking the activity of PDI with inhibitors and PDI blockade with an antibody reduced the endocytosis of KIR2DL4 and the amount of HLA-G bound to KIR2DL4.

**Figure 4:**
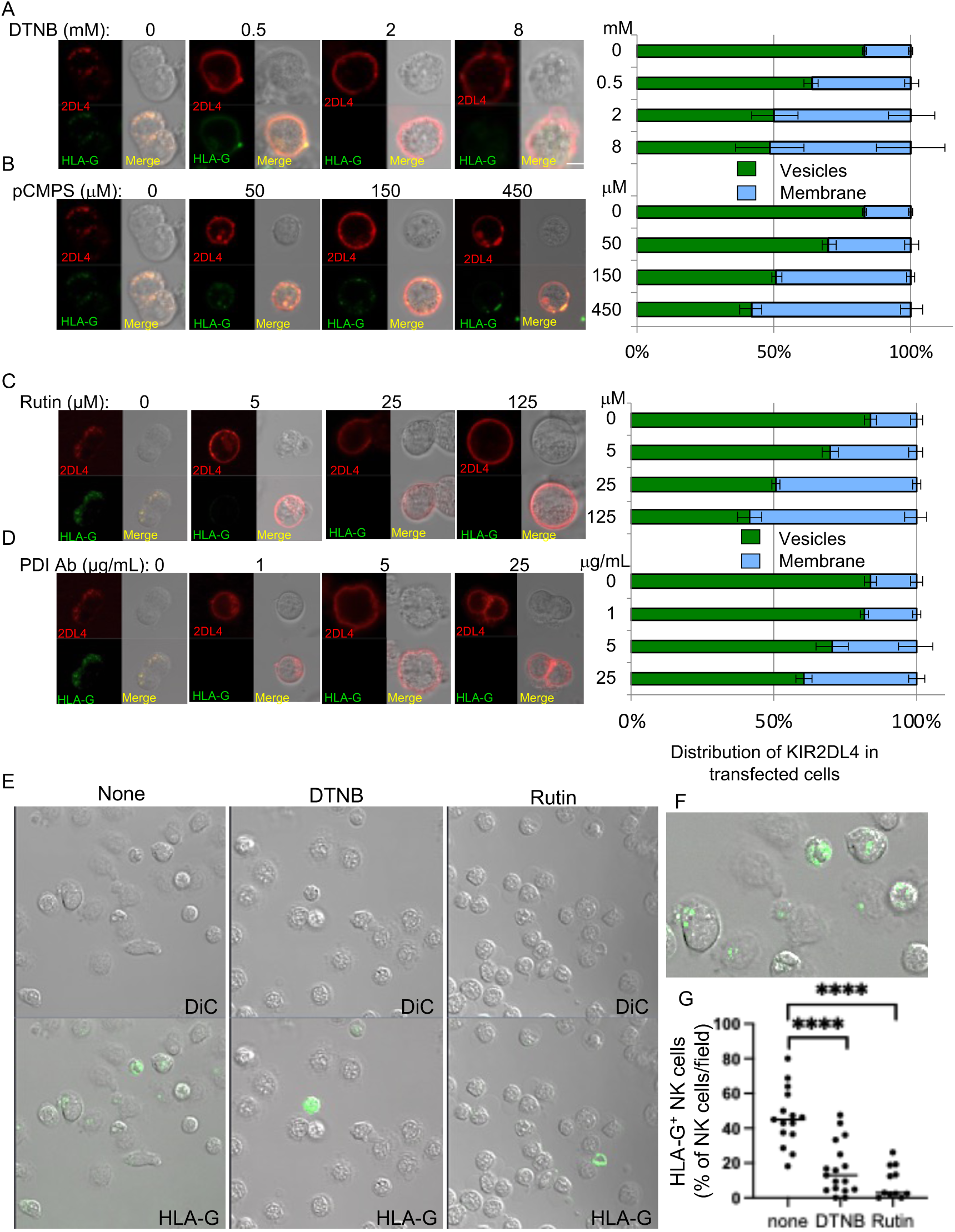
Location of KIR2DL4 and HLA-G after inhibition of PDI. (**A** to **D**) Confocal microscopy images of 293T cells transfected with HA-KIR2DL4 (anti-HA coupled to Alexa 594, *red*) and loaded with labeled HLA-G (*green*) for 2 h in the presence of DTNB (**A**), pCMPS (**B**), Rutin (**C**) and anti-PDI antibody (**D**), at the indicated concentrations. The time 0 image in (**A**) and (**B**) is the same, and so is the time 0 image shown in (**C**) and (**D**). The distribution of KIR2DL4 in vesicles or on the cell surface was determined by scoring at least 200 cells per condition. (**E**) Confocal microscopy images of primary resting human NK cells loaded with labeled HLA-G (green) for 2 h in the presence of 1 mM DTNB or 30 μM Rutin. The number of HLA-G positive cells were scored in multiple fields (11-18 fields/condition). (**F**) Enlarged image of Fig. 4E (bottom left) with vesicular localization of HLA-G. (**G**) Quantification of data obtained from analyzing experiments using NK cells from 3 different donors.

We then examined the impact of PDI on HLA-G loading by KIR2DL4 in primary NK cells. Resting, freshly isolated NK cells from three different donors were tested for the effect of PDI inhibition using the thiol blocker DTNB and the inhibitor Rutin at 30 μM, a concentration that is selective for PDI (*PDIA1*) (*23*). Cells were pretreated with inhibitors for 1 h prior to incubation with soluble HLA-G-Alexa-647 for 2 h in the continued presence of inhibitor and counted by confocal microscopy for cells that had endocytosed HLA-G (Fig. 4E). The fraction of NK cells containing a signal for HLA-G in vesicles (as shown in Fig. 4F) was counted in multiple fields for each condition (Fig. 4G). Treatment of cells with DTNB resulted in a significant decrease in intracellular HLA-G, as compared to untreated cells. Treatment with Rutin resulted in even stronger inhibition of soluble HLA-G loading (Fig. 4G). Thus, inhibition of PDI prevented HLA-G uptake by KIR2DL4 in both transfected 293T cells and in primary NK cells.

### Disulfide bonds in KIR2DL4 regulate HLA-G binding, endocytosis and function

To further evaluate the role of alternative disulfide bonds in the D0 domain and for a comparison of the same cysteine substitutions, each one of the three cysteines in the D0 domain was replaced with a serine. The C10S and the C28S mutants remained mostly at the plasma membrane (Fig. 5A), indicating that a Cys28-Cys74 configuration or a Cys10-Cys74 configuration (if it were to exist) did not internalize into vesicles. Conversely, the C74S mutant, which is limited to a Cys10-Cys28 bond, was distributed in intracellular vesicles like wild-type KIR2DL4 (Fig. 5A).

**Figure 5:**
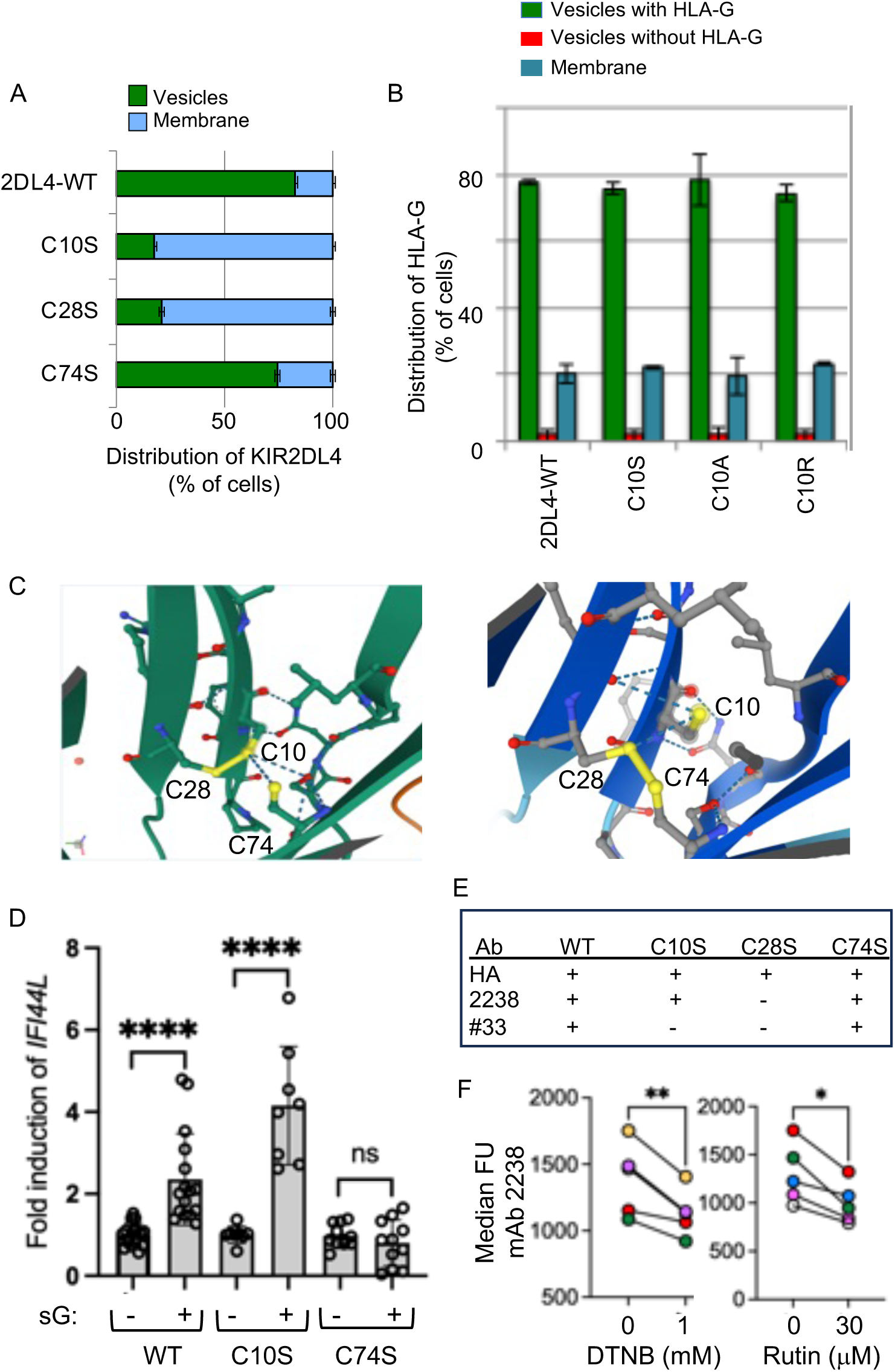
Cysteine mutants in KIR2DL4 tested for their impact on KIR2DL4 distribution. (**A**) Distribution (vesicles or plasma membrane) of WT and of the indicated cysteine mutants of KIR2DL4 in transfected 293T cells. (**B**) Distribution of HLA-G in 293T cells transfected with KIR2DL4 wild-type and with mutations at Cys10. HLA-G was scored for presence or absence in KIR2DL4+ vesicles. (**C**) Close-up view of the trio of cysteines in the D0 domain of KIR2DL4, as reported in the crystal structure (3WYR, *left*) and as predicted by AlphaFold (*right*). (**D**) Quantitative PCR analysis of *IFI44L* expression in response to HLA-G stimulation of 293T cells with stable expression of WT, C10S or C74S KIR2DL4. Baseline *IFI44L* transcription in untreated cells was set at 1. Data are represented as mean ± SD and are from 8 to 15 independent experiments depending on the cell line. ****p≤ 0.001; ns, p>0.05 by Wilcoxon test. (**E**) Reactivity of the antibodies anti-HA tag, mAb #33 and mAb 2238 with wild-type and the 3 cysteine mutants of KIR2DL4. (**F**) Flow cytometry analysis of primary NK cells pretreated with DTNB (1 mM) or Rutin (30 μM) for 1 h at 37°C and stained for surface expression of KIR2DL4 using mAb 2238 at 4°C in the presence of inhibitor for 30 min, followed by staining goat anti-mouse secondary antibodies coupled to APC at 4°C for 30 min. Data are represented as median fluorescence units and are from NK cells from 5 donors tested separately.

Considering that HLA-G bound to KIR2DL4 in a Cys28-Cys74 configuration, several mutants of Cys10 were tested for their ability to bind and internalize HLA-G. Three substitutions of Cys10 in HA-KIR2DL4 (C10S, C10A and C10R) transfected into 293T cells were able to bind HLA-G and to internalize as well as wild-type (Fig. 5B). There was very little internalization of HA-KIR2DL4 into vesicles without concomitant uptake of HLA-G (Fig. 5B), as was seen with wild-type KIR2DL4 (Fig. 1B). Therefore, KIR2DL4 with a Cys28-Cys74 disulfide bond behaved as wild-type, as far as HLA-G binding and transport to intracellular vesicles was concerned but failed to internalize on its own. A summary of cysteine mutants and their impact on KIR localization and interaction with HLA-G is shown in Table 1. Each one of the 3 cysteines were mutated to Ser, Ala, and Arg, which resulted in consistent phenotypes for C10 mutants (internalization induced by soluble HLA-G) and C28 mutants (no HLA-G binding, no internalization), whereas C74 mutants, which included also a Cys to Trp mutation, had either a clear-cut phenotype (internalization and no HLA-G binding with C74W and C74A), or an intermediate phenotype (some HLA-G internalization with C74S and C74R).

The X-ray crystal structure of a KIR2DL4 that had been produced and purified from human cells (*12*) revealed an unexpected cross-β strand Cys10-Cys28 bond in D0, rather than the canonical cross-β sheet Cys28-Cys74 (Fig. 5C). By comparison, AlphaFold 2, a deep learning-based prediction tool to model a protein 3D structure from its amino acid sequence (*24, 25*), predicted a KIR2DL4 structure with a Cys28-Cys74 bond (Fig. 5C), loosely supporting the notion that KIR2DL4 can exist in both configurations and consistent with an allosteric transition between two disulfide bonded states.

To determine which of the two configurations of KIR2DL4 is functional, we examined the ability of the C10S and the C74S mutants of KIR2DL4 expressed in 293T cells to generate functional outcomes in response to soluble HLA-G. Recently, single cell transcriptomics of primary NK cells stimulated with HLA-G revealed the induction of an ISG response in the absence of type I interferon (*10*). This response was recapitulated in HEK293 reporter cells transfected with KIR2DL4 (*10*). We generated 293T cells stably expressing HA-tagged wild-type (WT), C10S, and C74S KIR2DL4 with a comparable expression level (Fig. S3). Stimulation of these cells with soluble HLA-G for 16 h showed that *IFI44L* transcription was induced in cells expressing WT and C10S but not in cells expressing the C74S mutant (Fig. 5D). Therefore, a functional response to soluble HLA-G, such as ISG induction, is triggered by KIR2DL4 with a Cys28-Cys74 bond.

Our agonist mAb #33 and the commercial mAb MAB2238 (mAb 2238) to KIR2DL4 were tested for binding to KIR2DL4 carrying each of the three different Cys mutations. As control, an antibody to the HA-tag fused to the extracellular N-terminus of KIR2DL4 was used (Fig. 5E, Fig. S4). Neither mAb #33 nor mAb 2238 bound to the C28S mutant, presumably misfolded due to the lack of a Cys10-Cys74 disulfide bond, as shown by mass spectrometry (Fig. 3E). While they both bound to the C74S mutant, limited to a Cys10-Cys28 bond, only mAb 2238 bound to the C10S mutant, limited to a Cys28-Cys74 bond (Fig. 5E). Therefore, mAb #33 does not recognize KIR2DL4 in a Cys28-Cys74 configuration and binds selectively to the Cys10-Cys28 disulfide bonded state (Fig. 5E and Fig.S4).

We tested mAb #33 and mAb 2238 for their binding to KIR2DL4 on primary resting NK cells (Fig. S5). There was no detectable cell surface staining by mAb #33, indicating that the receptor in the Cys10-Cys28 configuration does not remain at the plasma membrane, but is rapidly endocytosed. This is consistent with our previous work showing that labelled antibody is endocytosed into early endosomes where the receptor resides (*14*). In contrast, upon staining with mAb 2238, which recognizes both disulfide forms, a subpopulation of primary NK cells expressed KIR2DL4 at the cell surface, presumably in the Cys28-Cys74 configuration. To test if cell surface levels of receptors in this configuration are sensitive to inhibition of PDI by DTNB and Rutin, we pretreated primary NK cells with inhibitors for 1 h, followed by staining for cell surface KIR2DL4 using mAb 2238 (Fig. 5F). Inhibition of PDI resulted in decreased surface staining of KIR2DL4 by mAb 2238. Since PDI targets the Cys10-Cys28 bond (Fig. 3F), our data is consistent with the interpretation that inhibition of PDI prevents the switch from Cys10-Cys28 to the Cys28-Cys74 form, detectable with mAb 2238 only.

### Impact of cysteine mutants on KIR2DL4 oligomerisation

KIR2DL4, unlike other members of the KIR family, forms dimers and higher-order oligomers in solution (*12*). Structural analysis of KIR2DL4 revealed distinct intermolecular interfaces that mediate both dimerisation and oligomerisation. Moreover, the presence of three internal cysteine residues within the KIR2DL4 D0 domain (Cys10, Cys28, Cys74) resulted in the formation of the non-canonical disulfide bond Cys10–Cys28 (instead of Cys28–Cys74) (*12*). This alternative disulfide bond arrangement was associated with conformational changes in D0 residues, which are widely conserved in KIR3D D0 domains, suggesting a potential role in regulating KIR2DL4 assembly (*12*). To investigate this, we tested the consequences of alternative disulfide bonding in the cysteine triad of KIR2DL4. To promote canonical or non-canonical disulfide bonding, respectively, we expressed and purified the KIR2DL4 point mutants C10L and C74S. Notably, both mutants were expressed at similar levels and purified via immobilised metal affinity chromatography (IMAC). However, while wild-type KIR2DL4 eluted as a mixture of oligomers during size exclusion chromatography, KIR2DL4(C10L) behaved as a KIR2DL4 dimer. In contrast, KIR2DL4(C74S) formed soluble aggregates (Fig. S6). These observations indicate that the cysteine triad within the KIR2DL4 D0 domain may promote higher-order oligomer formation.

### KIR2DL4 binds to the HLA-G**^VLPH^** epitope

To investigate HLA-G recognition by KIR2DL4, we conducted surface plasmon resonance (SPR) experiments to measure interactions between KIR2DL4 and HLA-G bound to peptide VLPHETRLL (HLA-G^VLPH^ epitope). While a previous study showed that KIR2DL4 did not interact with the epitopes HLA-G^RIIPRHLQL^ or HLA-G^RLPKDFRIL^ (*12*), we observed that KIR2DL4 bound to immobilised HLA-G^VLPH^ epitope with an equilibrium dissociation constant (K_D_) of 19.4 μM ± 4.00 μM (Fig. 6A) but did not interact with the HLA-G^VFPTKDVAL^ or HLA-G^RIIP^ epitope, the latter of which was consistent with our previous report (Fig. 6B). These data suggest that KIR2DL4 binds HLA-G epitopes in a peptide-specific manner with binding affinities comparable to those reported for KIR3DL1–HLA interactions (*20*). We subsequently investigated the ability of oligomerisation-deficient KIR2DL4(Cys10Leu) to recognise HLA-G^VLPH^ epitope and measured binding of KIR3DL1 as a negative control.

**Figure 6:**
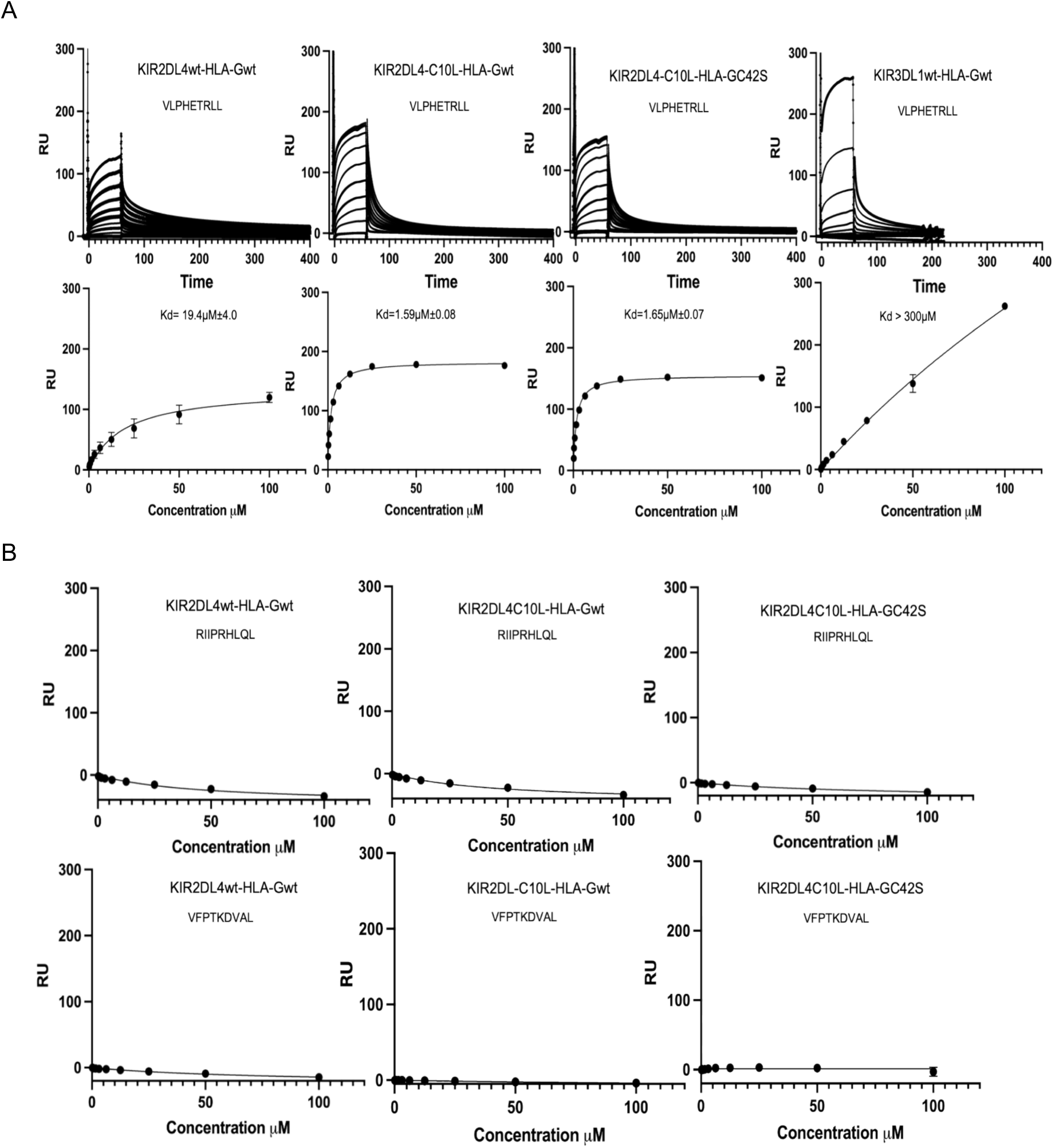
SPR affinity measurements. (A) Representative sensorgrams from single dilution series of KIR2DL4, KIR2DL4 (C10L) and KIR3DL1 binding to HLA-G^VPLH^ (top panel) and interaction analysis of this binding (bottom panel). (B) Analysis of KIR2DL4 and KIR2DL4(C10L) interaction with non-binding epitopes. Error bars denote standard deviation. Equilibrium dissociation constants K_D_ were determined using a 1:1 binding model from two independent experiments performed in duplicate.

Moreover, since native HLA-G forms disulfide-linked dimers, we further examined KIR2DL4(C10L) interactions with the monomeric C42S mutant HLA-G(C42S)^VLPH^ along with wild-type HLA-G (Fig. 6A). Our data showed that KIR2DL4(C10L) bound both wild-type HLA-G^VLPH^ and HLA-G(C42S)^VLPH^ with markedly higher affinity than wild-type KIR2DL4, with K_D_ values of 1.59 ± 0.08 μM and 1.65 ± 0.07 μM, respectively (Fig 6A). However, KIR2DL4(C10L) did not bind to HLA-G^VFPT^ or HLA-G^RIIP^, showing that KIR2DL4 recognises the HLA-G^VLPH^ epitope in a peptide-specific manner (Table S2). As previously shown, KIR2DL4 is expressed as a broad distribution of dimers and readily forms higher-order oligomers in a concentration-dependent manner. In contrast, our present data show that the KIR2DL4(C10L) mutant abrogated the formation of higher-order oligomers. Together with its enhanced affinity for HLA-G, these results suggest that oligomerisation of KIR2DL4 competes with ligand engagement and may regulate receptor availability for HLA-G recognition.

### HLA-G binding is controlled by an allosteric disulfide in the KIR2DL4 D0 domain

Wild-type KIR2DL4 is in the Cys10-Cys28 configuration in the crystal structure (*12*), whereas a structure predicted by AlphaFold 2 (*24, 25*) is in the Cys28-Cys74 disulfide-bonded form (Fig. 7A). When given a C74R mutant of KIR2DL4, AlphaFold predicted a structure similar to that of the crystal (Fig. S7A). An overlay of the PDB 3WYR structure (*purple*) and the AlphaFold prediction for wild-type (*blue*) showed that a loop between the β-strand C and the short α helix in D0 may be different (Fig. 7A, *blue arrow*, and Fig. 7B). The side chains of V45 and P46 would be shifted by 5.3 Å and 5.4 Å, respectively, between the two predicted structures (Fig. 7B). Predictions of KIR2DL4 docking with HLA-G by AlphaFold 3 suggested that the side chains of the V45 to P48 loop in the Cys10-Cys28 crystal structure may be too distant to make contacts with HLA-G (Fig. 7C), whereas an interaction between HLA-G and KIR2DL4 in the Cys28-Cys74 configuration is more likely (Fig. 7D). The structure predicted for a C10R mutant (limited to a Cys28-Cys74 bond) is similar to that obtained for wild-type KIRDL4 (Fig. S7B). Structures of KIR2DL4–HLA-G complexes obtained experimentally are required to determine how HLA-G distinguishes the D0 domain in the two alternative disulfide bond configurations.

**Figure 7:**
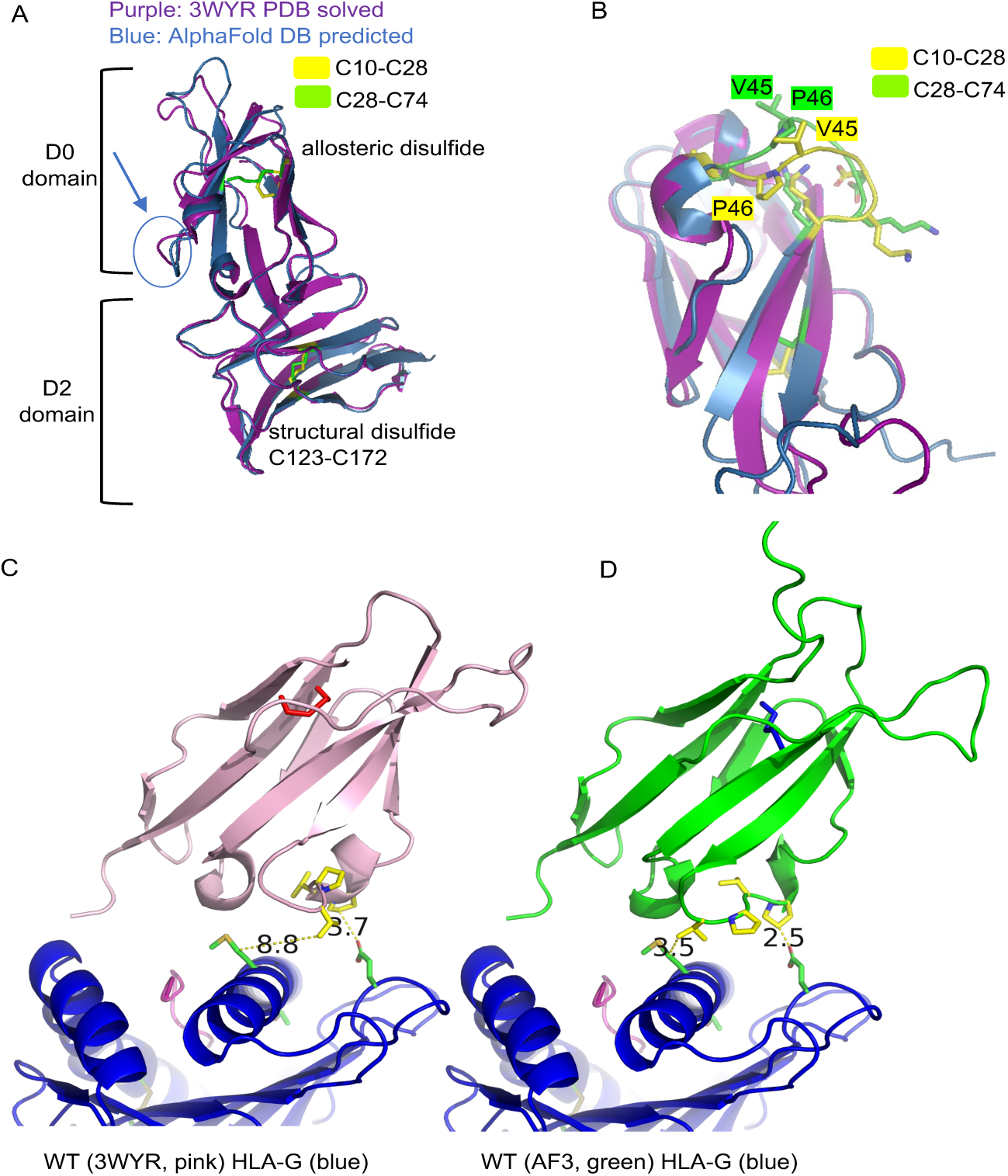
Predicted outcome of an allosteric disulfide switch on the conformation of KIR2DL4 and its predicted interaction with HLA-G. (**A**) Overlay of the crystal structure (3WYR; *purple*) and the structure predicted by AlphaFold DB (*blue*). The allosteric disulfide Cys10-Cys28 is in yellow and the Cys28-Cys74 structural disulfide is in green. The region indicated with a blue arrow corresponds to the loop highlighted in B. (**B**) D0 domain shown with the Pro46-Pro48 loop facing up and pointing toward the predicted HLA-G binding site. The position of Val45 and Pro46 in the crystal structure (C10-C28; *yellow*) is overlayed with that in the AlphaFold-predicted structure (C28-C74; *green*). The distance between Val45 and Pro46 in the crystal structure and their counterpart in the predicted structure of a Cys28-Cys74 KIR2DL4 is 5.3 and 5.4 Å, respectively. (**C**) Docking of the KIR2DL4 crystal structure (3WYR, *pink*) with HLA-G (*blue*), as predicted by AlphaFold 3. (**D**) Docking of the predicted KIR2DL4 structure by AlphaFold 3 (*green*) with HLA-G (*blue*). In the Cys28-Cys74 KIR2DL4 bonded form predicted by AlphaFold for wild-type KIR2DL4, the Val45-Pro48 loop appears to be in a different orientation, suggesting potential contacts with Met76 and Glu19 of HLA-G.

The conformation of the D2 domain is not predicted to be affected by the disulfide switch in D0, as seen in an overlay of D2 domains for all four structures, the crystal structure (3WYR) and the predicted structures of wild-type, C74R, and C10R KIR2DL4 (Fig. S6C). Thus, the D2 domain contacts may be preserved regardless of the disulfide bond used in the D0 domain. Taken together with the quantitative and direct binding data, these results support the interpretation that a distant allosteric change—induced by the switch from a Cys10-Cys28 disulfide bond to a canonical Cys28-Cys74 bond in KIR2DL4—is required for binding of HLA-G.

## Discussion

Transcription of *HLA-G* is turned on in fetal extravillous trophoblast (EVT) cells as they migrate to the fetal-maternal interface and invade the decidua during early pregnancy (*5*). The abundant maternal NK cells in the decidua are distinct from those in the blood (*26*) and have a unique transcriptional signature, including elevated transcription of *KIR2DL4* (*5*). As expected for cellular processes that support human reproduction, the encounter of EVT cells with decidual NK cells (dNK) at the maternal-fetal interface is well orchestrated (*5, 27*). High transcription of *KIR2DL4* in dNK cells is matched by high and selective transcription of *HLA-G* in EVT cells (*5*). EVT cells also upregulate transcription of several matrix metalloprotease genes (*5*), including *MMP2*, which promotes shedding of soluble HLA-G from the plasma membrane (*28, 29*). Uptake of soluble HLA-G into NK cells depends on KIR2DL4 and both colocalize in Rab5**^+^** early endosomes from where signaling occurs (*14, 30, 31*).

Transmembrane, full-length HLA-G at the cell surface is a ligand for the inhibitory receptor LILRB1 (*32, 33*), which is selectively expressed by the dNK1 subset of decidual NK cells (*5*). Thus, HLA-G at the plasma membrane serves to protect EVT cells from cytotoxic dNK1 cells. A second protective role of HLA-G occurs by the release of a peptide from its signal sequence, which binds HLA-E to form a strong ligand for the inhibitory receptor NKG2A-CD94 (*34–36*). Therefore, inhibitory signaling through LILRB1–HLA-G and NKG2A–HLA-E interactions together protect EVT cells from NK-mediated killing. At the same time, when EVT cells invade the decidua, soluble HLA-G shed from EVT cells by the activity of MMPs can be taken up by KIR2DL4 into endosomes of dNK cells, thereby bypassing the inhibition by LILRB1 at the plasma membrane. Soluble HLA-G promotes a secretory response in NK cells to support vascular remodeling, and a broad transcriptional response with a clear ISG signature (*4, 10*).

To investigate how KIR2DL4 interacts with HLA-G, a random mutagenesis approach was chosen to generate a library of KIR2DL4 mutants that were then tested for their ability to endocytose soluble HLA-G. This analysis revealed that binding and uptake of soluble HLA-G into endosomes was regulated by a trio of cysteine residues in the first Ig domain of KIR2DL4, called D0. In addition to the Cys28-Cys74 disulfide bond typical of Ig domains, such as the D0 domains of KIR2DL5 and KIR3DL1, a third cysteine is present at position 10 in KIR2DL4. The only KIR2DL4 structure reported so far has the atypical Cys10-Cys28 bond, leaving a free cysteine at position 74 (*12*). The purified receptor containing this bond did not bind HLA-G (*12*). KIR2DL4 mutations at Cys10, which leave the D0 domain with the Cys28-Cys74 bond as the only option, remained at the plasma membrane and internalized in response to soluble HLA-G. Thus, proper KIR2DL4 function depends on a switch from one disulfide bond to another (Cys10-Cys28 to Cys28-Cys74), keeping Cys28 as a partner for either Cys10 or Cys74.

We noticed that the Cys10-Cys28 bond in the KIR2DL4 crystal structure (*12*) has a – RHstaple conformation, which is a signature of allosteric bonds having an inherent high bond stress that predisposes them to cleavage (*22*). KIR2DL4 in human cells was found to exist in both disulfide configurations, and the allosteric Cys10-Cys28 bond was reduced in vitro by the thiol disulfide oxidoreductase PDI. As expected, if PDI activity is required to maintain a pool of Cys28-Cys74 KIR2DL4 at the plasma membrane (by reducing Cys10-Cys28 bonds), thiol blockade with membrane impermeable DTNB or inhibition of PDI with Rutin or a PDI-specific mAb did prevent HLA-G binding. AlphaFold predictions of KIR2DL4 folding suggest that a switch from Cys10-Cys28 to a Cys28-Cys74 disulfide bond may cause an allosteric change at a distant loop in D0 (Val45 to Pro48). In the Cys10-Cys28 configuration of KIR2DL4, the Val45 to Pro48 stretch may be oriented in a way not suited for HLA-G binding. Conversely, KIR2DL4 in the Cys28-Cys74 configuration is predicted to have the Val45 to Pro48 loop closer to HLA-G. The previously reported lack of binding of purified KIR2DL4 with HLA-G in vitro (*12*) may be explained by the incompatibility of a Cys10-Cys28 bonded KIR2DL4 with HLA-G binding.

Direct binding of HLA-G to KIR2DL4 was demonstrated by SPR with purified soluble KIR2DL4 and HLA-G associated with peptide VLPHETRLL. However, binding was undetectable with two other peptides tested, suggesting a strong dependence of KIR2DL4 on the sequence of the peptide used to refold HLA-G. Wild-type KIR2DL4 bound to HLA-G^VLPHETRLL^ with a K_D_ of 19 μM. Consistent with the KIR2DL4 mutagenesis study, stronger binding, with a K_D_ of 1.6 μM, was achieved with a purified Cys10Leu KIR2DL4 mutant (in a Cys28-Cys74 configuration). In addition, homo-dimerization of HLA-G via the free Cys42 was not required for binding to KIR2DL4, since a monomeric HLA-G with the Cys42Ser mutation that prevents dimer formation bound KIR2DL4 just as well as wild-type HLA-G.

Of interest, the peptide KGPPAALTL associated with HLA-G for our cellular assays was the most abundant peptide eluted from HLA-G from the placenta (*16*). It is derived from the protein encoded by *EBI3*, itself one of the strongly upregulated genes in invading EVT cells (*5*). A structure of HLA-G^KGPPAALTL^ showed a peptide bound in a linear manner with a relatively low solvent accessibility (*15*). Rules that dictate HLA-G binding to KIR2DL4 await further studies and structures of KIR2DL4–HLA-G complexes.

The regulation of a KIR2DL4–HLA-G interaction by an allosteric disulfide that controls protein conformation provides yet another example of disulfide bonds serving as switches in protein function. Allosteric disulfides regulate diverse biological processes, such as hemostasis (tissue factor F3, platelet fibrinogen receptor integrin αIIb-β3, and von Willebrand factor) and viral infection (HIV-1 gp120) (*18*). Allosteric disulfide bonds specifically targeted by extracellular PDI at the cell surface have been identified in proteins such as GP1bα, tissue factor (*F3*), integrin αvβ3 during thrombosis, and HIV-1 envelope gp120 when bound to CD4 during viral entry (*37, 38*).

*KIR2DL4*, the only KIR gene family member expressed by all human NK cell subsets, with a high transcription rate in dNK cells, joins the list of proteins whose function is controlled by a labile disulfide bond acting as a switch, but here in the context of early pregnancy. In response to fetal HLA-G, NK cells endocytose soluble HLA-G bound to KIR2DL4, which signals from endosomes for a secretory response that leads to a sustained secretion of factors to support vascular remodeling and fetal development (*4, 17*). In addition, NK cells mount an ISG transcriptional response upon endocytosis of HLA-G, which could protect them from viral infection during pregnancy (*10*). In this unique environment at the maternal-fetal interface, the allosteric disulfide bond regulates KIR2DL4 localization, binding to HLA-G, and signaling to enable the supportive role of NK cells for a successful pregnancy.

## Materials and Methods

### Cells and reagents

HEK293T cells (here called 293T) were obtained from the American Type Culture Collection (ATCC) and cultured in Iscoves Modified Dulbecco’s medium (IMDM) containing 10% fetal calf serum (FCS). KIR2DL4*001 with a single polymorphism in residue Glu101 to Gly (GenBank accession number DQ266438) was used in this study (unless otherwise indicated). KIR2DL4 and receptor 2B4 (*CD244*) were tagged at the N-terminus with the HA-tag peptide YPYDVPDYA present in the pDisplay expression vector. 293T–2DL4-gfp cells were generated as described (*14*). Briefly, 293T–2DL4-gfp cells were made by transfecting 293T cells using LipofectAMINE 2000, with a KIR2DL4 cDNA fused to green fluorescence protein (GFP) in the vector pBABE (puro) CMV^+^ using SalI and BamH1 sites. Transfected cells were selected and maintained in IMDM containing puromycin at 2 μg/mL. Primary NK cells from healthy human donors were isolated as follows: Peripheral blood samples from deidentified healthy US adults were obtained from the NIH Department of Transfusion Medicine under an NIH Institutional Review Board-approved protocol (99-CC-0168) with informed consent. Peripheral blood mononuclear cells (PBMC) were isolated by separating lymphocytes with Lymphocyte Separation Medium (MP Biomedical), after which non-NK cells were removed using the NK cell isolation kit (STEMCELL Technologies) supplemented with anti-CD3 beads. The purity of isolated resting NK cells was determined by flow cytometry. Cells that were > 95% CD3^-^, CD14^-^, CD16^+^, CD56^+^, CD69^-^, ILT2^-^ and NKp46^+^ were used. Primary NK cells were stimulated with HLA-G (2 μM) for 16 h at 37°C. Recombinant HLA-G and HLA-E were obtained from the Immune Monitoring Services, Fred Hutchinson Cancer Center (Seattle, WA). Briefly, MHC heavy chain protein and β2-microglobulin (β2m) were produced in *E. coli* and purified from inclusion bodies. The HLA-G heavy chain and β2m proteins were refolded *in vitro* in the presence of peptide KGPPAALTL into a monomeric peptide-MHC complex. For HLA-E, refolding was done in the presence of peptide VMAPRTLFL. Monomers of the refolded p-MHC were purified by FPLC size exclusion chromatography and concentrated to 1 mg/mL. Soluble HLA-G and HLA-E were labeled using the Alexa Fluor 647 Protein Labeling Kit (Cat #: A20173, Molecular Probes, ThermoFisher). mAb #33 was produced in our laboratory (*13*) and MAB2238 was obtained from RnD Systems.

### Random Mutagenesis Screen

Random mutagenesis of the KIR2DL4 sequence spanning amino acids 1-208, consisting of the Ig domains D0 and D2 and a portion of the stem was carried out using the Diversify PCR Random Mutagenesis kit (Clontech, Cat#630703) according to manufacturer’s instructions. HA-tagged KIR2DL4 expressed in pDisplay served as the DNA template. The primers used at 10 mM for PCR were as follows: Sense: CCTGCTATGGGTACTGCTGCTCTGGGTTCC; Antisense: GAAGGGGAGGATGGTGAAGAGGATGATGGC. Mn2**^+^** at 80 mM and dGTP at 40 mM were used to obtain the desired mutation rate of 1 to 2 mutations per kb of DNA. Thermal cycling conditions consisted of denaturation at 94° for 30 sec, followed by annealing at 94°C for 30 sec and extension at 68°C for 1 min for a total of 25 cycles. Mutated KIR2DL4 was cloned back into pDisplay, and clones were sequenced to select for single or double amino acid substitutions and remove frame shifts and stop codons. One μg of each mutant plasmid was transiently transfected into 293T cells using LipofectAMINE 2000 according to manufacturer’s instructions. After 36 h, cells were trypsinized and plated at a density of 7 × 10^4^ cells/well in eight-well coverglass slides (Nunc Lab-Tek, Thermo Fisher Scientific) 18 h prior to loading with anti-HA antibody or HLA-G. Cells were incubated with 0.03 mM anti-HA Ab coupled to Alexa 594 (Biolegend) and 0.04 mM HLA-G coupled to Alexa 647 for 2 h at 37°C. Cells were then washed in PBS and viewed by microscopy using a Zeiss 780 confocal microscope. The majority of mutants behaved like the wild-type KIR2DL4. Every mutant with a distinct phenotype was tested again, as above, after a new transfection, to confirm their phenotype.

### Confocal microscopy

Image acquisition was performed using a LSM 780 confocal laser scanning microscope (Zeiss) fitted with a LD C-Apochromat 40×/1.1 W Corr objective. Cells were incubated at 37°C in 5% CO_2_ for the entire image acquisition duration. The HFT 543 nm and 633 nm main beam splitters were used to detect signals from fluorophores Alexa-Fluor 594 and 647, respectively. Parameters were set to acquire at a fixed pixel density of 512 × 512, 16-bit pixel depth, and pinhole size of 100 μm. Images were processed, and colocalization coefficients were calculated using Zeiss ZEN software. Quantification of receptor and ligand localization was performed as follows. In transfected 293T cells expressing fluorescently tagged proteins, cells were viewed in a series of multiple fields using the LSM 780. Cells in each field were counted and categorized based on the location and ligand colocalization status of the expressed receptor. Counts were averaged over 2 to 4 experiments, and the number of cells counted per sample ranged from 100 ≤ n ≤ 430. In a typical transfection with wild-type KIR2DL4, approximately 80% of the transfected cells had clear signals for internalized HLA-G and KIR2DL4 in the same intracellular, punctate compartments, and the remaining 20% had low but detectable 2DL4 at the plasma membrane but no HLA-G. Primary NK cells, loaded with recombinant soluble HLA-G labelled with Alexa 647 for 2 h were viewed in multiple fields and HLA-G positive NK cells were counted as a fraction of all the NK cells in the field. The percentage of HLA-G loaded cells/field was calculated using NK cells from 3 different donors tested separately.

### Thiol Blockade and PDI Inhibition Assays

In eight-well coverglass slides, 293T cells transfected with HA-2DL4 were preincubated in IMDM containing DTNB (5,5′-Dithiobis 2-nitrobenzoic acid; Sigma), pCMPS (para-chloromercuriphenylsulfonate; Carbosynth), Rutin (Sigma), or anti-PDI antibody RL90 (Thermo Fisher) at varying concentrations for 1 h at 37°C. Anti-HA-594 and HLA-G-647 were then added and incubated for 2 h in the continued presence of inhibitor at 37°C. Cells were then washed twice with PBS and analyzed by confocal microscopy as described above. Primary NK cells were placed in eight well coverglass slides and preincubated in IMDM containing 1 mM DTNB or 30 μM Rutin for 1 h at 37°C. HLA-G-Alexa 647 was then added and incubated for 2 h in the continued presence of inhibitor at 37°C. Cells were then washed twice with PBS and analyzed by confocal microscopy as described above. Primary NK cells were also tested by flow cytometry for the effect of DTNB and Rutin (pretreated as described above for 1 h at 37°C) on surface expression of KIR2DL4 by staining with primary antibodies mAb #33 and mAb 2238 at 4°C, followed by staining with secondary goat anti-mouse antibodies coupled to Alexa 488 at 4°C. Stained cells were analyzed on a Cytoflex flow cytometer (Beckman).

### Quantification of cysteine redox states

GFP-tagged KIR2DL4 expressed in 293T cells was immunoprecipitated from cell lysates using ChromTek-GFP Trap Agarose (Proteintech). Free thiols were labelled with 2-iodo-N-phenylacetamide (^12^C-IPA) followed by cell lysis, immunoprecipitation of 2DL4-GFP and a second ^12^C-IPA labelling of proteins prior to elution. 2DL4-GFP immunoprecipitated from cells was analyzed on SDS-PAGE. Proteins within the size range of 2DL4-GFP (75 kDa) were excised from the gel area, reduced with dithiothreitol (DTT), and reduced thiols were alkylated with ^13^C-IPA. The redox states of cysteines were quantified by mass spectrometry analysis. Reactions were stopped by adding 5% (v/v) formic acid, and peptides were eluted from the gel slices with 5% formic acid and 50% (v/v) acetonitrile. Peptides were desalted using Ziptip (Millipore), dried and reconstituted in 0.1% formic acid. Using a Thermo Fisher Scientific Ultimate 3000, peptides were injected and resolved on a 35cm × 75 mm C18 reverse phase analytical column with integrated emitter using a 2-35% acetonitrile over 22 min with a flow rate of 300 nL/min. The peptides were ionized by electrospray ionization at +2.0 kV. Tandem mass spectrometry analysis was carried out on a Q-Exactive Plus mass spectrometer (Thermo Fisher Scientific) using Higher-energy Collisional Dissociation (HCD) fragmentation. The data-dependent acquisition method acquired MS/MS spectra of the top 5 most abundant ions at any one point during the gradient. The data was searched using Mascot (Matrix Science) against Uniprot database. Search parameters were as follows: precursor tolerance of 6 ppm and product ion tolerances of ± 0.4 Da. Oxidized Met, pyro-Glu and pyro-Gln were selected as variable modifications with full chymotryptic cleavage of up to three missed cleavages. To calculate ion abundance of peptides, extracted ion chromatograms were generated using the XCalibur Qual Browser software (v2.1.0; Thermo Scientific). The area was calculated using the automated peak detection function built into the software. Abundance of disulfide-linked peptides was expressed as a ratio relative to disulfide-linked peptide between Cys123-Cys172 in the D2 domain. Disulfide bond analysis was performed using the Disulfide bond Analysis Tool: (http://52.142.47.78/disulfideanalysis/).

Recombinant KIR2DL4 protein (WT, C10S and C74S) were produced in insect cells and purified by His-tag affinity chromatography. The extracellular domain of KIR2DL4*001 (residues 1-195) was subcloned into the pAC insect expression vector in frame with an N-terminal 8× HisTag and a 3C protease cleavage site. The KIR2DL4 expression vector was then transfected with BestBac 2.0 linearized DNA (Expression Systems) into Sf9 cells to generate P1 virus. Virus was amplified by shaking culture in Sf9 cells prior to expression. KIR2DL4 was then expressed in BTI-Tn-5B1-4 insect cells (High Five) and purified using Ni-NTA affinity purification followed by Superdex 200 size exclusion into 10 mM HEPES, 150 mM NaCl, pH 7.2. Fractions containing the purified protein were verified by SDS-PAGE. Recombinant thiol isomerases PDI (*PDIA1*), ERp57 (*PDIA3*) and thioredoxin (*TXN*) were expressed in *E. coli* and purified by His-tag affinity chromatography. The thiol isomerases were then reduced with final 10 mM dithiothreitol for 30 min at 25°C and desalted into PBS using a Zeba spin column (Thermo Fisher Scientific). KIR2DL4 (3 mg) was treated with either a 2- or a 10-fold molar excess of each thiol isomerase for 30 min at 25°C. Free thiols in 2DL4 were alkylated with 5 mM 2-iodo-N-phenylacetamide (^12^C-IPA) and the protein was resolved on non-reducing SDS-PAGE, stained with Bio-Safe Coomassie stain and destained in deionized water. Bands corresponding to 2DL4 were excised, dried, incubated with 40 mM dithiothreitol and washed. The fully reduced proteins were alkylated with 5 mM ^13^C-IPA. The gel slices were washed and dried before digestion of proteins with 12 ng/mL of chymotrypsin (Roche) in 25 mM NH_4_CO_2_ containing 10 mM CaCl_2_ overnight at 25°C. Peptides were eluted from the slices with 5% formic acid and 50% acetonitrile. Peptides were desalted using Ziptips (Merck Millipore). Liquid chromatography, mass spectrometry and data analysis were performed as described (*39, 40*). The fraction of reduced disulfide bond was measured from the relative ion abundance of peptides containing ^12^C-IPA and ^13^C-IPA. To calculate ion abundance of peptides, extracted ion chromatograms were generated using the XCalibur Qual Browser software (v2.1.0; Thermo Fisher Scientific). The area was calculated using the automated peak detection function built into the software. The ratio of ^12^C-IPA and ^13^C-IPA alkylation represents the fraction of the cysteine in the population that is in the reduced state (*39*).

### Site directed mutagenesis

Engineered mutations of KIR2DL4 in the pDisplay vector were carried out using the QuikChange site directed mutagenesis kit (Stratagene) according to the manufacturer’s instructions and verified by sequencing. Stable transfections of cysteine mutants of KIR2DL4 were carried out using Lipofectamine 2000 according to the manufacturer’s instructions and selected using 1 μg/mL puromycin. Transfected cells were sorted by flow cytometry to select transfectants with uniform expression of the HA-tag.

### q-PCR analysis

Total RNA was extracted using RNeasy^®^ Plus Micro Kit from QIAGEN (Cat#74034). 1 μg of total RNA was used to prepare cDNA using iScriptTM cDNA Synthesis Kit (BioRad Cat# 1708891). Each RT-q-PCR assay was done in triplicate with 10 ng of cDNA/assay and detected in CFX Connect Real-Time System from BioRad using the manufacturer’s recommended standard thermal cycling protocol. Briefly, polymerase activation and DNA denaturation was done at 95°C for 3 min, denaturation at 95°C for 10 seconds, and annealing/extension at 55°C for 30 seconds. Amplification of *IFI44L* gene was done for 39 cycles. *ACTB* was used as the housekeeping reference gene. Relative expression was calculated using 2^(average housekeeping gene – average experimental gene) and results are presented as fold induction (HLA-G-stimulated/untreated control).

### Protein production and Surface Plasmon Resonance measurements

DNA sequences encoding KIR2DL4*001, KIR2DL4*001(Cys10Leu) and KIR3DL1*001 and an N-terminal His6-Tag were cloned into the pHLsec vector and expressed in Human Embryonic Kidney (HEK) Expi293F cells (GIBCO). Culture supernatants were dialysed against HBS500 buffer (10 mM HEPES pH 7.5, 500 mM NaCl) and proteins were subsequently purified via IMAC and size exclusion chromatography as previously described (*12, 20*). Wild-type and Cys42Ser mutant of HLA-G were produced and purified as previously described (*41*). Briefly, wild-type HLA-G and HLA-G(Cys42Ser) alpha chain with C-terminal BirA biotinylation signal, as well as human β2M, were separately expressed and isolated from *E. coli* as inclusion bodies. HLA-G complexes were obtained by refolding HLA-G BirA alpha chains, β2M, along with CMV pp65 protein epitopes pp65-2 (VLPHETRLL) and pp65-6 (VFPTKDVAL) (*42*), and H2A peptide (RIIPRHLQL) (*41*) under the denaturing conditions at 4°C for two days. Proteins were dialysed against 10 mM Tris, pH 8.0 and purified using DEAE and Hitrap-Q ion-exchange chromatography. Purified HLA-G complexes were concentrated and biotinylated using BirA ligase and further purified via S200 size-exclusion chromatography. SPR measurements were performed on a BIAcore T200 instrument (GE Healthcare) at 25°C in 10 mM HEPES (pH 7.5), 300 mM NaCl, 0.005% surfactant P20, as described (*43*). Wild-type and Cys42Ser HLAG-BirA loaded with pp65-VLPHETRLL epitope and HLA-A2^S269-277^ as negative control were immobilised at a flow rate of 5 μL/min on a streptavidin sensor chip (Cytiva). Binding interactions were determined by passing serial dilutions of KIR protein with a maximum concentration of 100 μM over immobilised HLA complexes at a flow rate of 10 μl/min for 60s, followed by 300s and 120s dissociation time for wild-type KIR2DL4 and KIR2DL4 (Cys10Leu) or KIR3DL1, respectively. Two independent experiments with two replicates were performed for each KIR, and equilibrium dissociation constants KD were determined by fitting a single-site binding model to the data.

## Acknowledgements

We thank the Department of Transfusion Medicine at NIH for blood samples and Joseph Brzostowski at the Twinbrook Imaging Center (NIAID) for assistance with confocal microscopy. This research was supported by grant ZIA AI000525 (E.O.L.) from the Intramural Research Program (IRP) of the National Institutes of Health (NIH). It was also supported by grant RO1 AI170952 (E.J.A) from NIAID, NIH; the National Health and Medical Research Council of Australia grants 2008981 (J.R.) and 2026631 (P.J.H.); a Senior Researcher Grant from the NSW Cardiovascular Research Capacity Program (P.J.H.); the Heart Foundation Vanguard Grant 110585-2025 (J.C.), and St George and Sutherland Research Medical Foundation Standard Grant RG254054 (J.C.). The contributions of the NIH authors were made as part of their official duties as NIH federal employees, are in compliance with agency policy requirements, and are considered Works of the United States Government. However, the findings and conclusions presented in this paper are those of the authors and do not necessarily reflect the views of the NIH or the U.S. Department of Health and Human Services.

## Supplementary information for

**Fig. S1:**
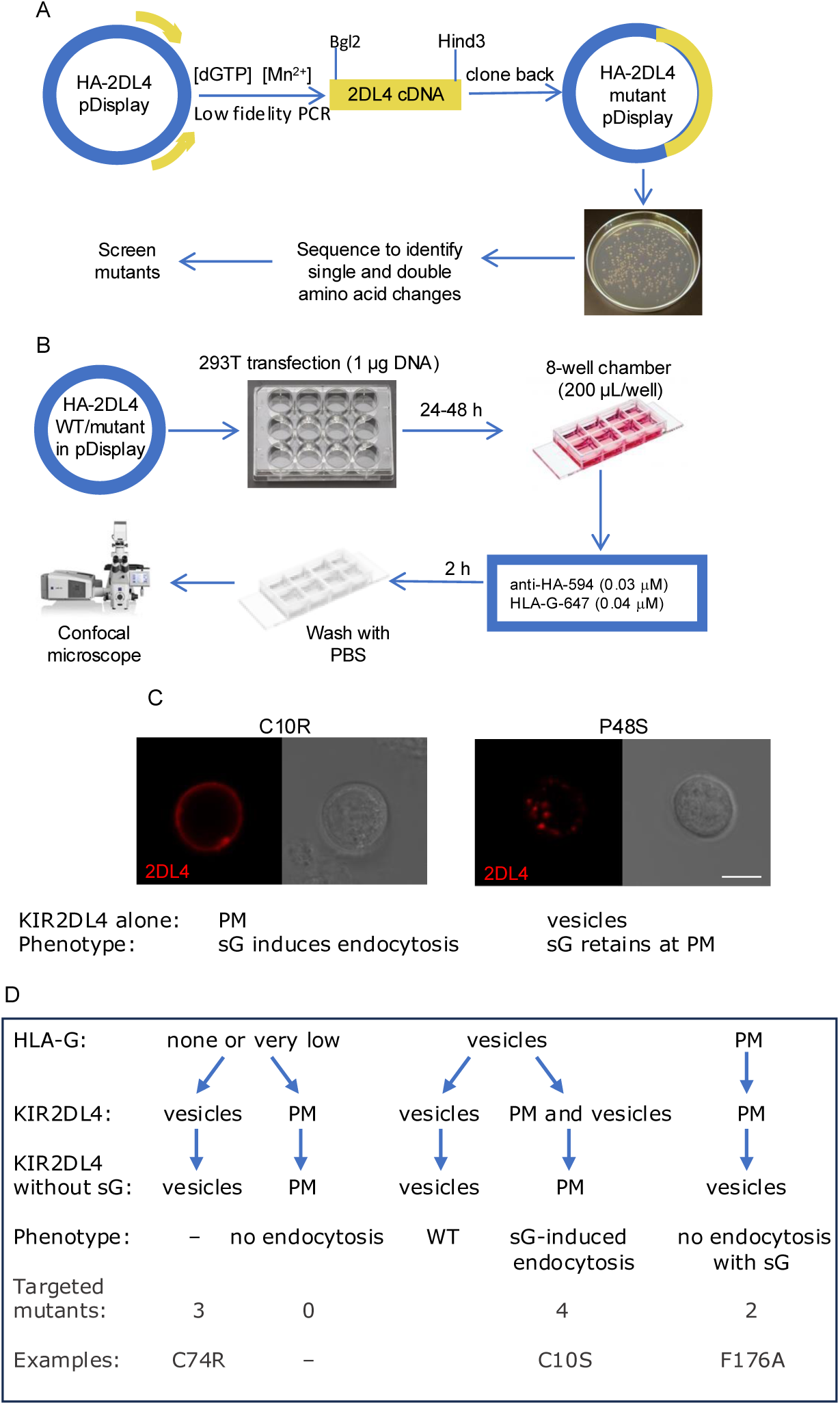
Random mutagenesis screen. (**A**) Strategy for the generation of mutants. (**B**) Examination of mutants by confocal microscopy. WT = wild-type. (**C**) In the same assay as in Figure 1C, mutants C10R and P48S were tested in the absence of soluble HLA-G (sG). Confocal images of anti-HA-Alexa-594 revealed their location. A DIC image is shown in the right panel. PM = plasma membrane. (**D**) Flow chart of mutants distributed among 4 categories distinct from WT. Several mutants from each category were also tested in the absence of soluble HLA-G (sG). HLA-G often dictates the location of KIR2DL4. For example, C10R internalization depended on HLA-G, and P48S was not internalized into vesicles in the presence of soluble HLA-G.

**Fig. S2:**
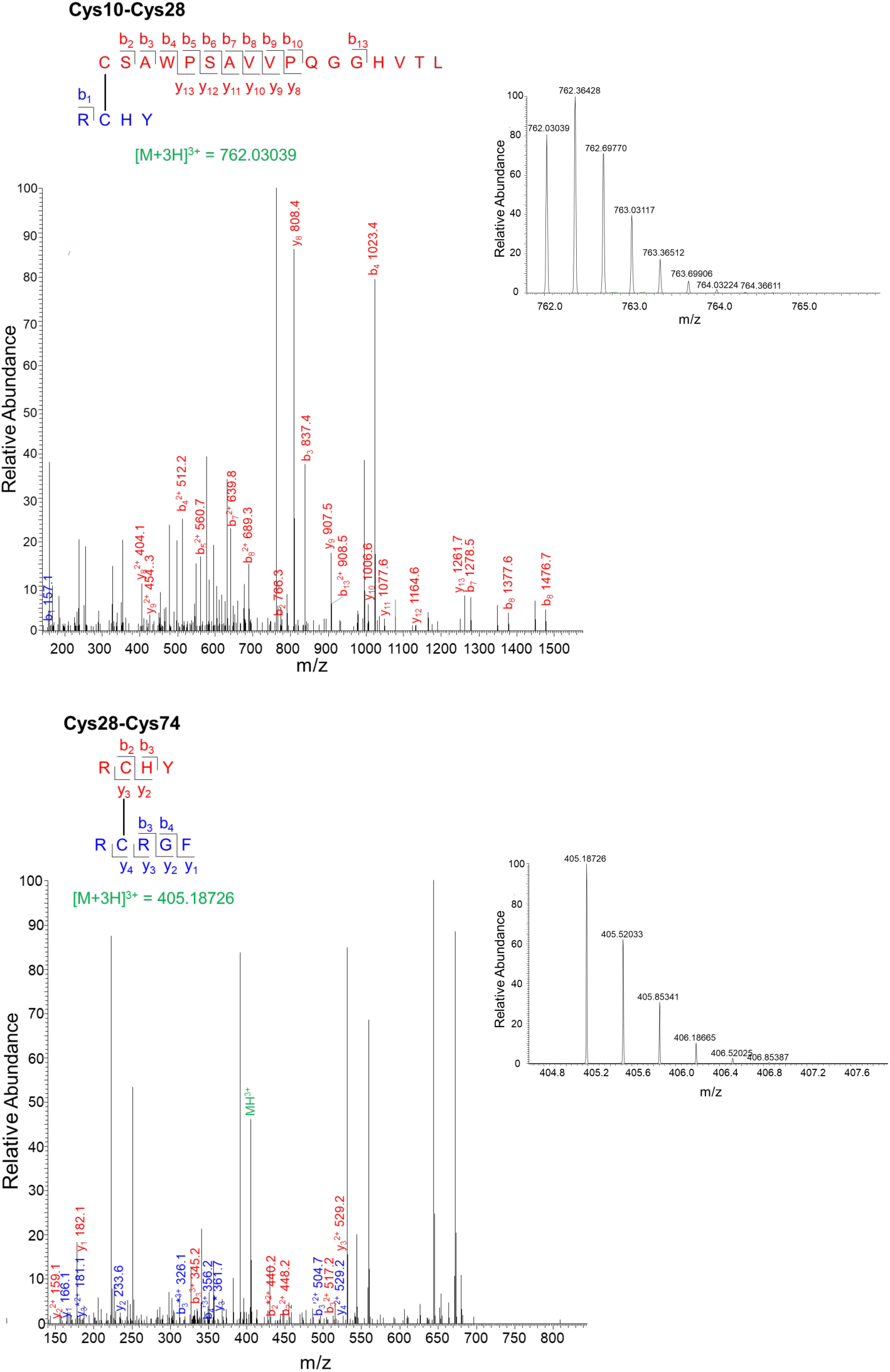
Analysis of disulfide-linked peptides in recombinant KIR2DL4 protein by mass spectrometry. Tandem mass spectra of the peptides linked by disulfide bond between Cys10 and Cys28 and between Cys28 and Cys74 are shown. The accurate mass of Cys10-Cys28 and Cys28-C74 peptides are shown in the insets (Cys10-Cys28: [M + 3H]3+ = *m*/*z* 762.03039 and expected [M + 3H]3+ = *m*/*z* 762.0299; Cys28-Cys74 : [M + 3H]3+ = *m*/*z* 405.18726 and expected [M + 3H]3+ = *m*/*z* 405.1870).

**Fig. S3:**
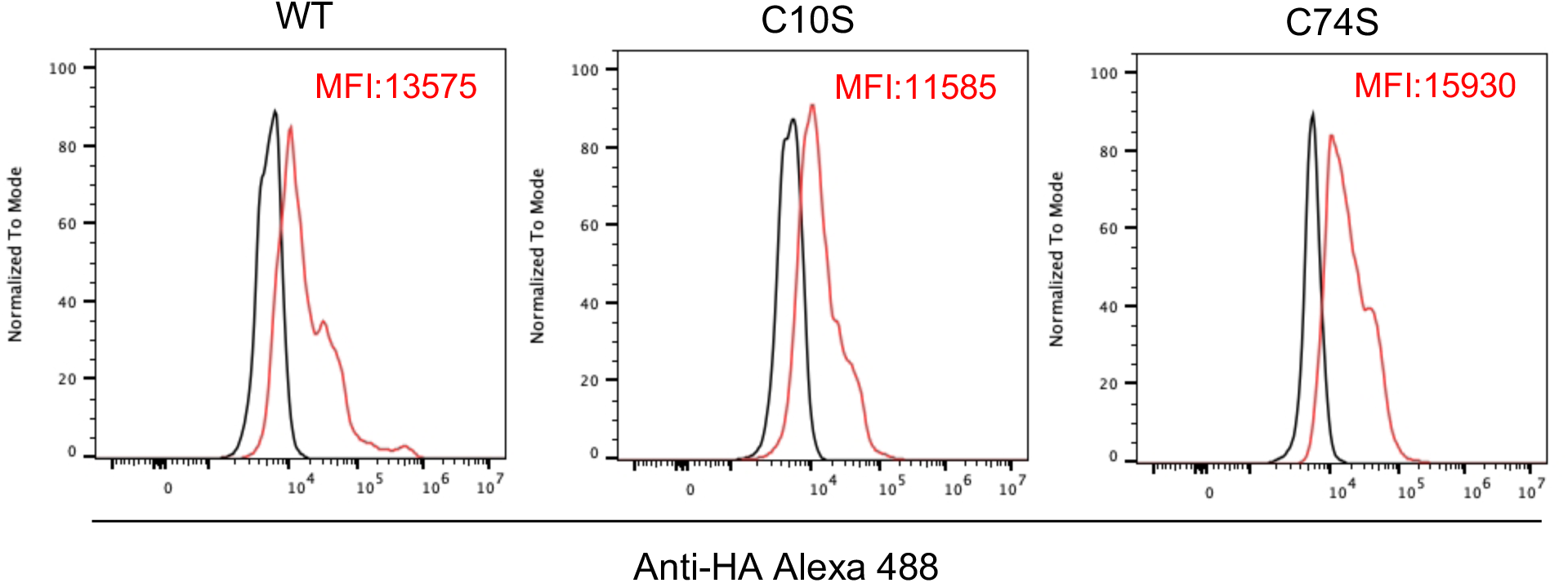
Flow cytometry profiles of 293T cells stably expressing WT, C10S or C74S mutants of HA-tagged KIR2DL4. Cells were stained with anti-HA tag antibody coupled to Alexa 488. Median fluorescence intensity (MFI) for the red profiles is indicated in the upper right corner.

**Fig. S4.**
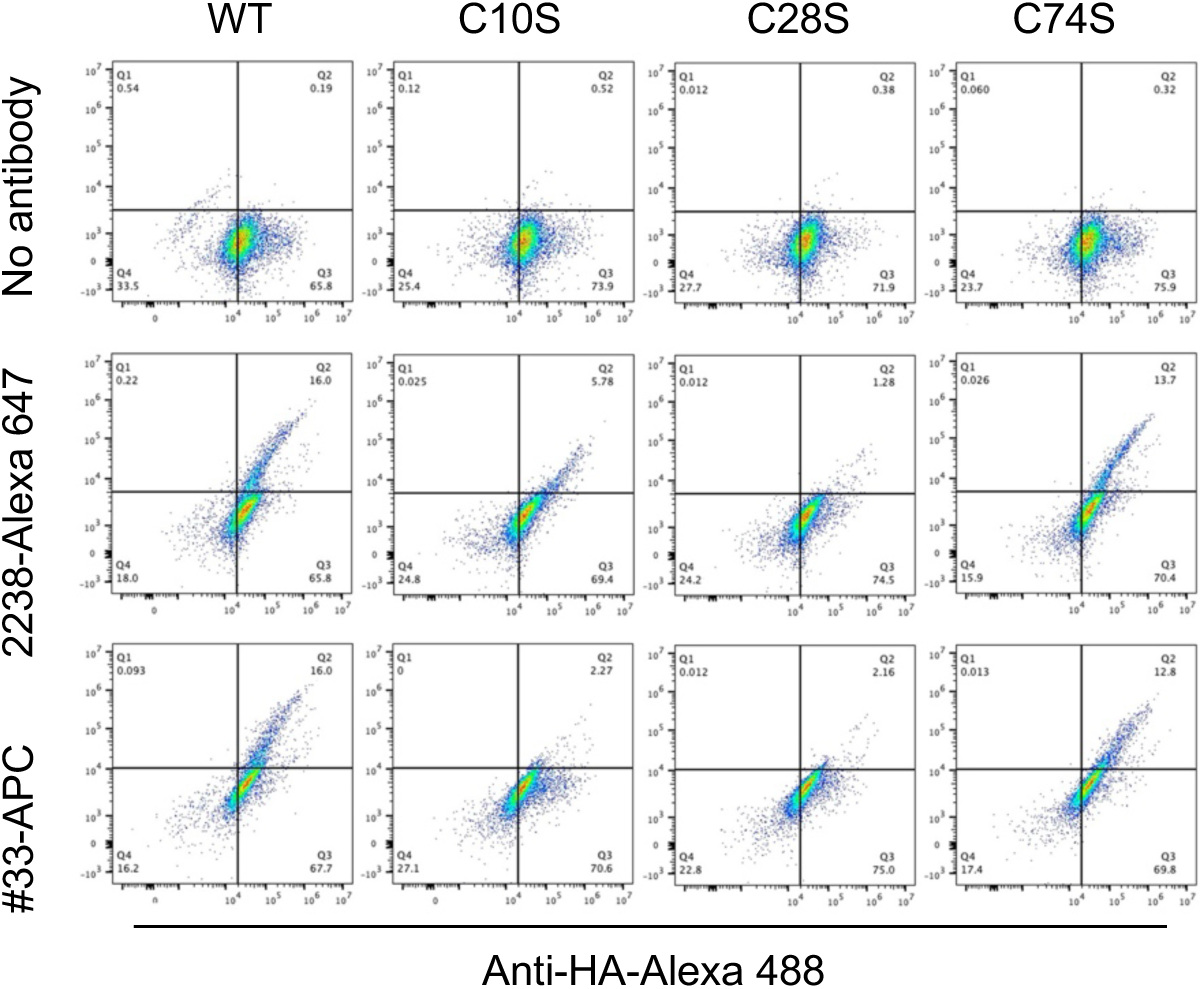
Flow cytometry profiles of 293T cells expressing HA-tagged KIR2DL4 WT and, in parallel, the 3 Cys to Ser mutants loaded for 2 h at 37°C with an antibody to the HA-tag and two different antibodies to KIR2DL4. Staining with the control anti-HA antibody coupled with Alexa 488 is shown on the x axis versus no antibody (*top*), whereas mAb 2238-Alexa 647 (*center*) and mAb #33-APC (*bottom*) are on the y axis. The lower left quadrant was set to include 98% of unstained control cells.

**Fig. S5.**
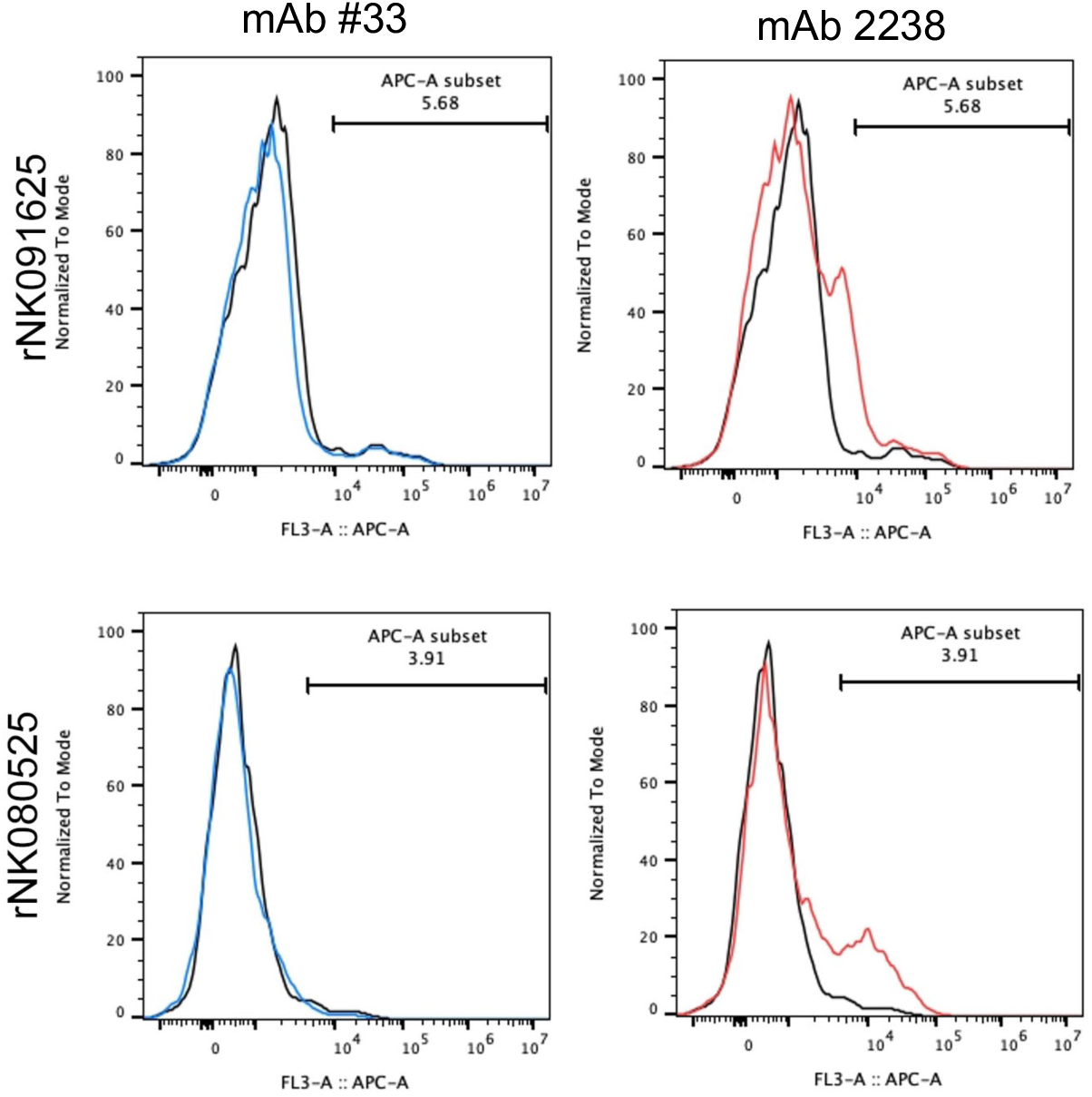
Flow cytometry profiles of primary NK cells from two different donors stained at the cell surface with mAb #33 (blue profiles) or mAb 2238 (red profiles) followed by APC-coupled secondary antibodies. Control staining with secondary antibodies alone is also shown (black profiles).

**Fig. S6.**
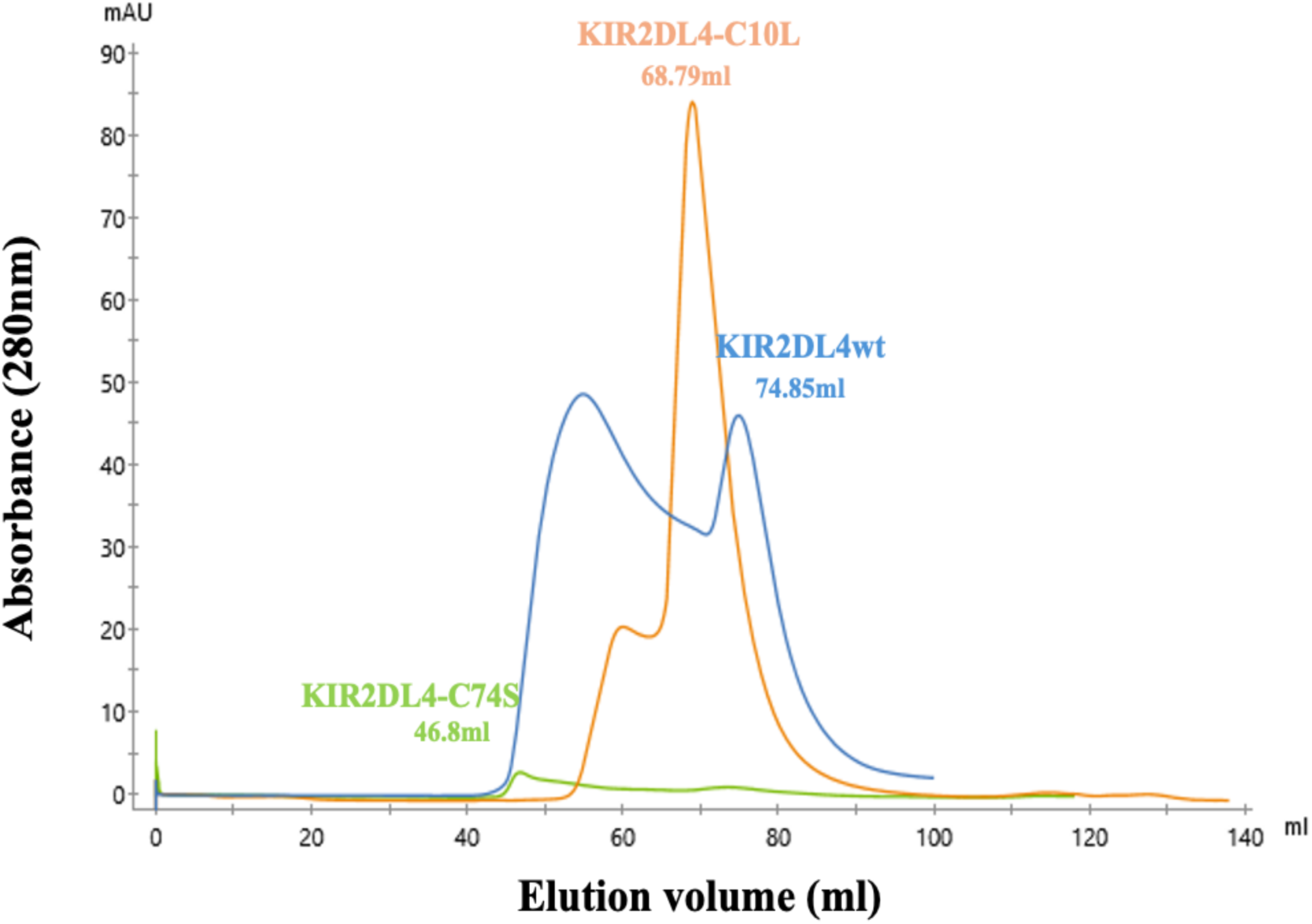
Size exclusion chromatography (SEC) profiles. Overlay of size exclusion profile with elution volume of purified KIR2DL4 wt (blue), KIR2DL4 C10L (orange) and KIR2DL4 C74S (green).

**Fig. S7.**
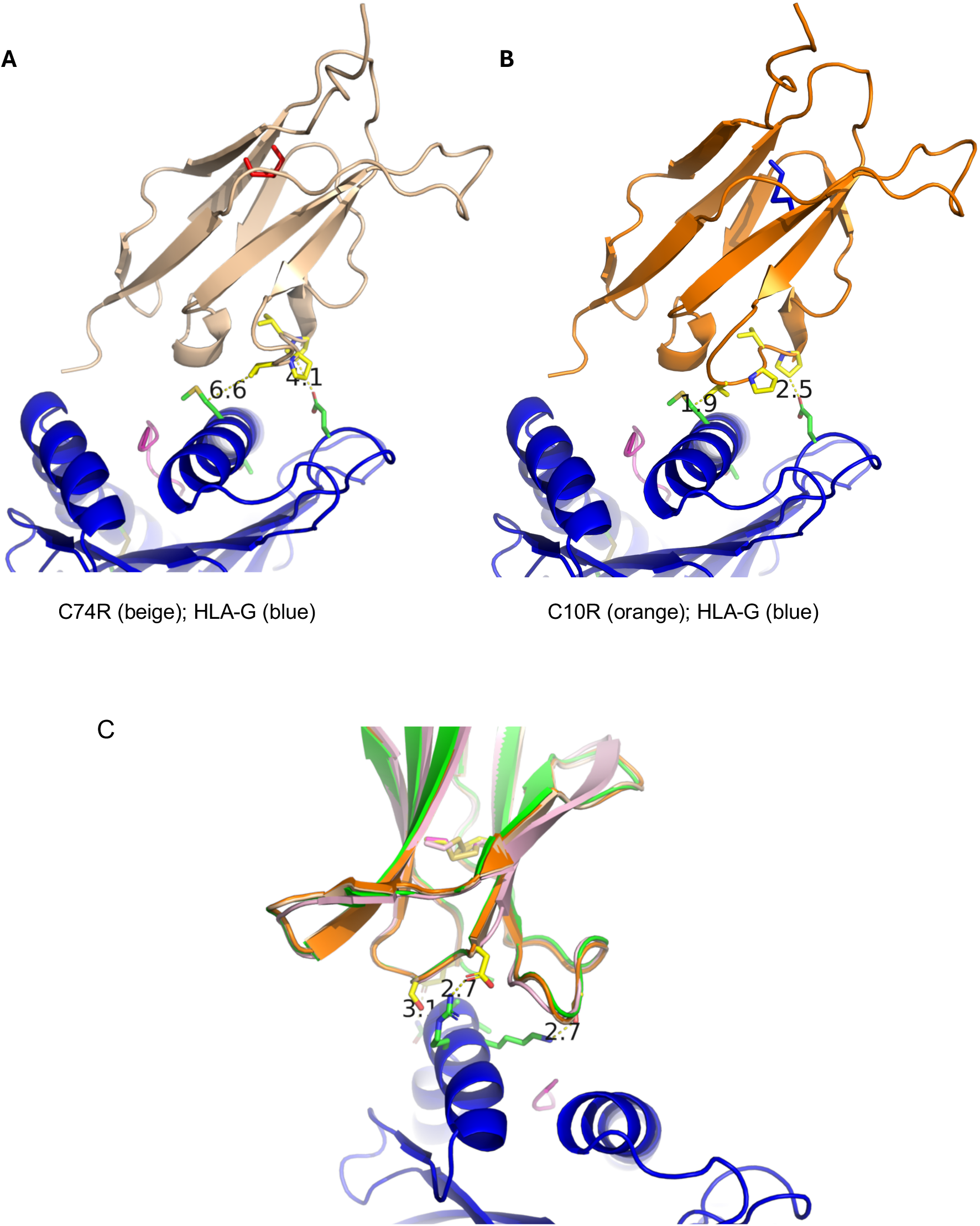
Potential interaction of KIR2DL4 with HLA-G, as predicted by AlphaFold (**A**) Docking of the C74R KIR2DL4 (*beige*) with HLA-G (*blue*), as predicted by AlphaFold 3. The structure of KIR2DL4 in this Cys10-Cys28 configuration is similar to that of the crystal structure (Fig. 7C). (**B**) C10R KIR2DL4 (*orange*) with HLA-G (*blue*). The predicted structure of this Cys28-Cys74 KIR2DL4, with the Val45-Pro48 loop closer to HLA-G, resembles that of WT KIR2DL4 (Fig. 7D). (**C**) Predicted folding of the D2 domain in the four KIR2DL4 isoforms (3WYR crystal structure of WT, and AlphaFold predictions of WT, C74R, and C10R) examined here, indicate a conserved D2 domain folding (shown as an overlay). Several predicted contacts of KIR2DL4 with HLA-G (*blue*) are similar to those of KIR3DL1 D2 domain with HLA-B, such as Ser128–Arg145 (3.1 Å, *left*), Asp130–Arg145 (2.7 Å, *center*), and Ser179–Lys146 (2.7 Å, *right*). As KIR2DL4 lacks the 100 amino acid-long D1 domain of KIR3DL1, the Ser128, Asp130, and Ser179 of KIR2DL4 correspond to Ser228, Asp230, and Ser279 of KIR3DL1.

**Table S1.** KIR2DL4 peptides analysed to determine disulfide bond redox state. Peptides were detected by Byonic analysis software, confirmed by MS/MS and have errors <6 ppm. Only peptides with peak areas >10 million for a given Cys were included in the analysis.

| **Cys** | **Peptide sequence\*** | **Peptide score<sup>#</sup>** | **Peptide error (ppm)** |
| --- | --- | --- | --- |
| 10 | **CSAWPSAVVPQGGH**VTL | 549.9 | 0.16 |
|  | **CSAWPSAVVPQGGH** | 403.3 | 0.8 |
|  | **CSAW** | 351.1 | 0.8 |
| 28 | **RCHY** | 255.5 | 0.3 |
| 74 | **RCRGF** | 245.7 | 0.72 |
| 123 | **SCSSQSSF** | 383.9 | 0.97 |
|  | **SCSSQSSFDIY** | 387.2 | 0.32 |
| 172 | **RCFGSF** | 367.9 | 0.91 |

\*Cys was labelled with <sup>12</sup>C-iodoacetanilide or <sup>13</sup>C-iodoacetanilide and is in bold.
<sup>#</sup>Peptide score calculated by Byonic analysis software is a primary indicator of peptide-spectrum matches.

**Table S2.** Summary of SPR measurements.

| Epitope | HLA | KIRs | K<sub>D</sub> (μM) |
| --- | --- | --- | --- |
| Pp65-2-VLPHETRLL | HLA-G wt | KIR2DL4*001 | 19.4 μM ± 4.00 |
| Pp65-2-VLPHETRLL | HLA-G wt | KIR2DL4*001C10L | 1.59 ± 0.08 |
| Pp65-2-VLPHETRLL | HLA-GC42S | KIR2DL4*001C10L | 1.65 ± 0.07 |
| Pp65-2-VLPHETRLL | HLA-G wt | KIR3DL1*001 | >300 |
| Pp65-2-VLPHETRLL | HLA-GC42S | KIR3DL1*001 | >300 |
| H2A-RIIPRHLQL | HLA-G wt | KIR2DL4*001 | N.B. |
| Pp65-6-VFPTKDVAL | HLA-G wt | KIR2DL4*001 | N.B. |
| H2A-RIIPRHLQL | HLA-G wt | KIR2DL4*001C10L | N.B. |
| Pp65-6-VFPTKDVAL | HLA-G wt | KIR2DL4*001C10L | N.B. |
| H2A-RIIPRHLQL | HLA-GC42S | KIR2DL4*001C10L | N.B. |
| Pp65-6-VFPTKDVAL | HLA-GC42S | KIR2DL4*001C10L | N.B. |

KIR binding to the immobilised HLA-G epitopes was measured in two independent experiments. K<sub>D</sub> was calculated from all data using one site-specific binding model. N.B. No binding.

## Notes

### Competing Interest Statement

The authors have declared no competing interest.

### Summary of Updates

New authors: Priyanka Chaurasia, Jan Petersen, and Jamie Rossjohn, at Monash University, in Clayton, Victoria 3800, Australia. They contributed new data that demonstrate direct binding of HLA-G to KIR2DL4. New experiments with primary NK cells to confirm data obtained with transfected cells. New experiments to show that the transcriptional response by KIR2DL4 to stimulated with HLA-G occurs with one but not the other disulfide-bonded form of KIR2DL4.

